# Xiphoid nucleus of the midline thalamus controls cold-induced food seeking

**DOI:** 10.1101/2023.03.16.533067

**Authors:** Neeraj K. Lal, Phuong Le, Samarth Aggarwal, Alan Zhang, Kristina Wang, Tianbo Qi, Zhengyuan Pang, Dong Yang, Victoria Nudell, Gene W. Yeo, Alexander S. Banks, Li Ye

**Affiliations:** Department of Neuroscience, The Scripps Research Institute, La Jolla, CA 92037, USA; Department of Molecular Medicine, The Scripps Research Institute, La Jolla, CA 92037, USA; Department of Cellular and Molecular Medicine, University of California San Diego, La Jolla, CA, USA; Division of Endocrinology, Diabetes and Metabolism, Beth Israel Deaconess Medical Center, Boston, MA, USA

## Abstract

Maintaining body temperature is calorically expensive for endothermic animals. Mammals eat more in the cold to compensate for energy expenditure, but the neural mechanism underlying this coupling is not well understood. Through behavioral and metabolic analyses, we found that mice dynamically switch between energy conservation and food-seeking states in the cold, the latter of which is primarily driven by energy expenditure rather than the sensation of cold. To identify the neural mechanisms underlying cold-induced food seeking, we use whole-brain cFos mapping and found that the xiphoid (Xi), a small nucleus in the midline thalamus, was selectively activated by prolonged cold associated with elevated energy expenditure but not with acute cold exposure. *In vivo* calcium imaging showed that Xi activity correlates with food-seeking episodes in cold conditions. Using activity-dependent viral strategies, we found that optogenetic and chemogenetic stimulation of cold-activated Xi neurons recapitulated cold-induced feeding, whereas their inhibition suppressed it. Mechanistically, Xi encodes a context-dependent valence switch promoting food-seeking behaviors in cold but not warm conditions. Furthermore, these behaviors are mediated by a Xi to nucleus accumbens projection. Our results establish Xi as a key region for controlling cold-induced feeding, an important mechanism for maintaining energy homeostasis in endothermic animals.

---

The emergence of endothermy brought numerous adaptive advantages during evolution; however, it also came with a significant increase in energy expenditure. To fuel this increased energy demand, mammals dramatically adapt their foraging behavior in response to changing temperature, and there is a tight inextricable association between ambient temperature and food intake: the colder the environment, the more food is needed to maintain core body temperature^1–3^. Mammals, including humans, are known to eat more in the cold. Various festivals across different cultures with lavish feasts during winters are a testimony to our endothermic evolutionary past^4–6^. However, the neural basis linking the energy needs arising from the cold and the increase in feeding remains one of the unanswered questions in our understanding of mammalian biology.

Rodents are an excellent model to study this association between temperature and energy consumption^7^. For example, laboratory mice (*Mus musculus*) can double their daily food intake when living at 4°C, with thermogenesis contributing to ∼60% of the whole-body energy expenditure in these conditions^8^. Although cold sensation has been shown to acutely influence feeding through conserved somatosensory and feeding centers in the brain in both ectotherms and endotherms^9–13^, it remains unclear whether or how cold-induced energy expenditure is compensated by food intake. Here, we combined high-resolution metabolic and behavioral analyses to demonstrate that the increased feeding in the cold is a consequence of energy expenditure; moreover, we identified the midline thalamic Xiphoid nucleus (Xi) as a key hub mediating the compensatory increase of food-seeking behaviors.

## Cold-induced feeding is a result of elevated energy expenditure

Housing mice at 4°C has been reported to lead to elevated thermogenesis, energy expenditure (EE), and food intake^8,14^. To gain a refined view of EE, we used indirect calorimetry to determine an individual mouse’s EE in real-time through the measurement of oxygen and carbon dioxide exchanges^15^. Temporally resolved indirect calorimetry showed that switching environment temperature from 23°C to 4°C immediately increased EE (Fig. 1a-b, Extended Data Fig.1a-f). However, there was an initial decrease and substantial delay between the temperature drop and increased food intake (Fig. 1b), suggesting that the rapid cold sensation might not be the direct cause of cold induced feeding. Upon quantification of the temporal association between EE and feeding, we found that as cold exposure progressed, food intake became more correlated with EE (starting between 5-6 hours post-cold exposure and remaining elevated thereafter, Fig.1c). Based this progressive correlation, we thus defined this 5–6-hour post-cold exposure period as the onset of the cold-induced energy compensation (CIEC) and focused on this time window for the rest of the study, including video recorded mice within this hour to better understand these behaviors.

**Fig. 1.**
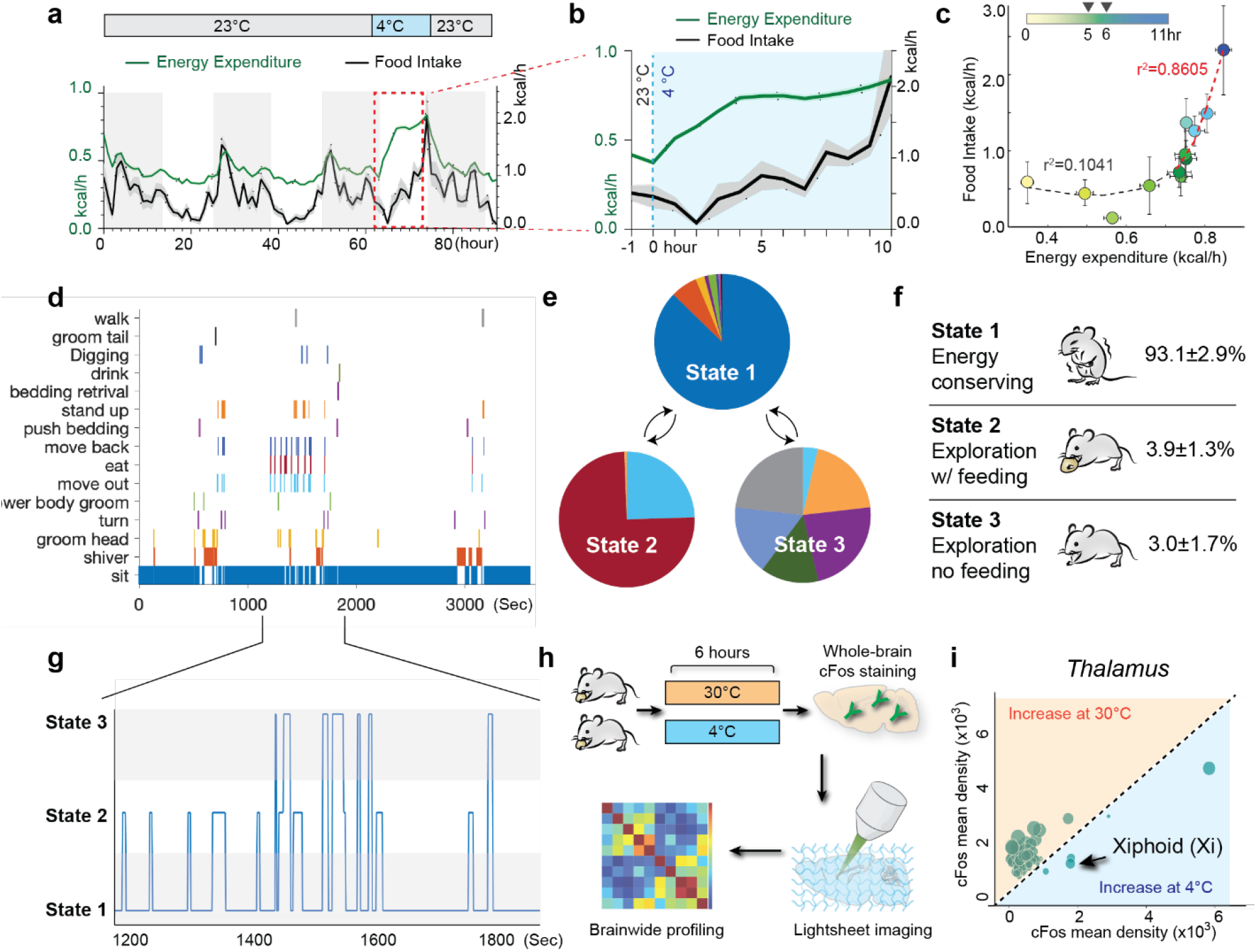
Identification of the behavioral and circuit basis of cold-induced feeding. **a**, 90-hour progression of energy expenditure (EE) and food intake (n=24 mice). Mice were housed at 23°C for 62 hours before switching the housing temperature to 4°C for 11 hours then reverting to 23°C. EE is charted in green and food intake in black; lines and shading denote the mean and SEM of each group. **b**, Zoomed-in from (a) during the switch from 23°C to 4°C. **c**, Scatter plot representing the relationship between the EE and feeding over an 11-hour period post-temperature switch (during the light cycle, n=24 mice). Each dot represents the average EE and food intake across all mice at each hour. Locally weighted scatterplot smoothing (LOWESS) was implemented to generate a lowess curve showing the r2. The error bars represent mean ± SEM both horizontally and vertically. Light yellow dots are hours closer to the onset of cold, whereas the dark green dots are later hours after the onset of cold. The two black arrows indicate the key transition period between 5-6 hours after onset of cold. **d-f**, Behavior analysis of animals undergoing cold-induced energy compensation (CIEC) using Hidden Markov Model (HMM). Manually annotated behavior videos (180 minutes long) from mice undergoing CIEC was used to generate the input sequence. The input sequence of each mouse was put into a vector. We defined a 3-state HMM: (1) energy-conserving state, (2) exploration with feeding, and (3) exploration without feeding. We generated the initial guess for the estimated transition matrix by assuming equal probabilities for all transitions as we did not have a priori information. **d**, Representative behavioral events assessed for a male C57BL/6J mouse after being in the cold for 5 hours. **e**, Representation of the HMM of CIEC-associated feeding (n=3 mice). Using labeled sequences of behaviors from **d**, the HMM estimated emission matrix of three states. **f**, Percentage of time spent in each HMM state. **g**, Zoomed-in from a 10-min session of **e**, showing the HMM state assignment. **h**, Schematics of whole-brain clearing and volumetric 3D imaging used to identify brain regions activated during CIEC. **i**, cFos mapping results for the thalamus. Each dot represents the cFos+ cell count in each distinct region based on Allen Brain Atlas registration. The size of the dots represents the difference in cell counts between the two conditions. Structures activated in thermoneutral temperature are shaded in orange and those activated in the cold are shaded in blue. Besides the Xi, four other thalamic structures were activated by the cold: Paraventricular nucleus of the thalamus (PVT), Intermediodorsal nucleus of the thalamus (IMD), Interanteromedial nucleus of the thalamus (IAM), and Midline group of the dorsal thalamus (MTN).

Interestingly, despite the overall increase in food intake when experiencing cold, the mice mostly stayed immobile rather than actively engaging in foraging (Fig. 1d). This observation suggested that instead of a unilateral increase in appetite, the mice faced competing priorities between conserving energy for thermogenesis (by staying immobile) versus replenishing energy supply (by seeking food). However, mice also engaged in various non-feeding, thermoregulatory behaviors in the cold, such as bedding retrieval and arrangement (Fig. 1d). To better understand the relationships among these behaviors, we first annotated them into 15 specific actions and used an unsupervised Hidden Markov Model (HMM) to identify three distinct states displayed by the mice: state 1, an energy conserving state in which the mice were mostly immobile; state 2, an exploration with food-seeking state with the highest probability of eating; and state 3, an exploration without food-seeking state, where mice engaged in other actions without food consumption (Fig. 1d-g and Extended Data Fig. 2a-c). This analysis enabled us to use just three HMM states, rather than 15 specific actions, to selectively study the CIEC-induced feeding state (state 2). As state 1 is the predominant state (Fig1. f), we focused on characterizing outbound state transitions from state 1 for all subsequent analyses.

### A brain-wide screen identified cold-induced activation of the Xi nucleus

To identify the circuit mechanisms underlying CIEC-induced feeding, we first performed a brain-wide cFos screen^16,17^ from mice that had been in the 4°C or 30°C (murine thermoneutrality, for maximizing brain-wide activity differences) for 6 hours, using whole-brain SHIELD, immunofluorescence labeling, lightsheet imaging, and automated analysis (Fig. 1h, Supplementary Table 1) ^18–20^. Cold conditions led to a broad decrease of cFos signal in the cortex, likely due to the overall decrease in physical activity of these animals (Extended Data Fig. 3a-c). The hypothalamus, where thermoregulation is centered^21^, showed robust cFos activation as expected (Extended Data Fig. 3a and d). Surprisingly, while most of the thalamus was suppressed in cold, several ventral midline thalamic (vMT) regions showed higher activation in the cold (Fig. 1i). We decided to further explore this area for two reasons. First, in Siberian hamsters, a model for cold adaptations, lesions at the vMT have been reported to impair cold adaptation in winter^22^. Second, it was recently shown in mice that the vMT is a crucial hub for gating how internal states are translated into opposite behavior responses (e.g., freezing vs. fighting) to perceived threats^21^, a scenario which resembles the competing priorities between energy conservation and food-seeking behaviors we observed in CIEC.

Next, to differentiate if vMT activation was because of cold sensation or related to CIEC, we examined cFos expression after 5 hours (to model CIEC) or 15 minutes (to model acute sensation) of cold exposure (Fig. 2a). Both conditions induced cFos in the median pre-optic nucleus (MnPO), a key region for thermoregulation (Extended Data Fig 4b). However, only 5 hours of cold exposure led to cFos activation in the Xiphoid nucleus (Xi) compared to 15-minute cold exposure or housing at 30°C; this activation was not observed in the neighboring Re (Fig. 2b-c, Extended Data Fig. 4a). Following our designation of 5–6-hour post-cold exposure period the onset of CIEC (Fig. 1c), we define these cFos-positive Xi neurons as the Xi^CIEC^ population, whose activation is likely due to CIEC rather than acute cold sensation. This selective Xi activation was also observed after an extended period of cold (7 days) (Extended Data Fig. 4c). Furthermore, both male and female mice showed activation of Xi during CIEC (Extended Data Fig 4d).

**Fig. 2.**
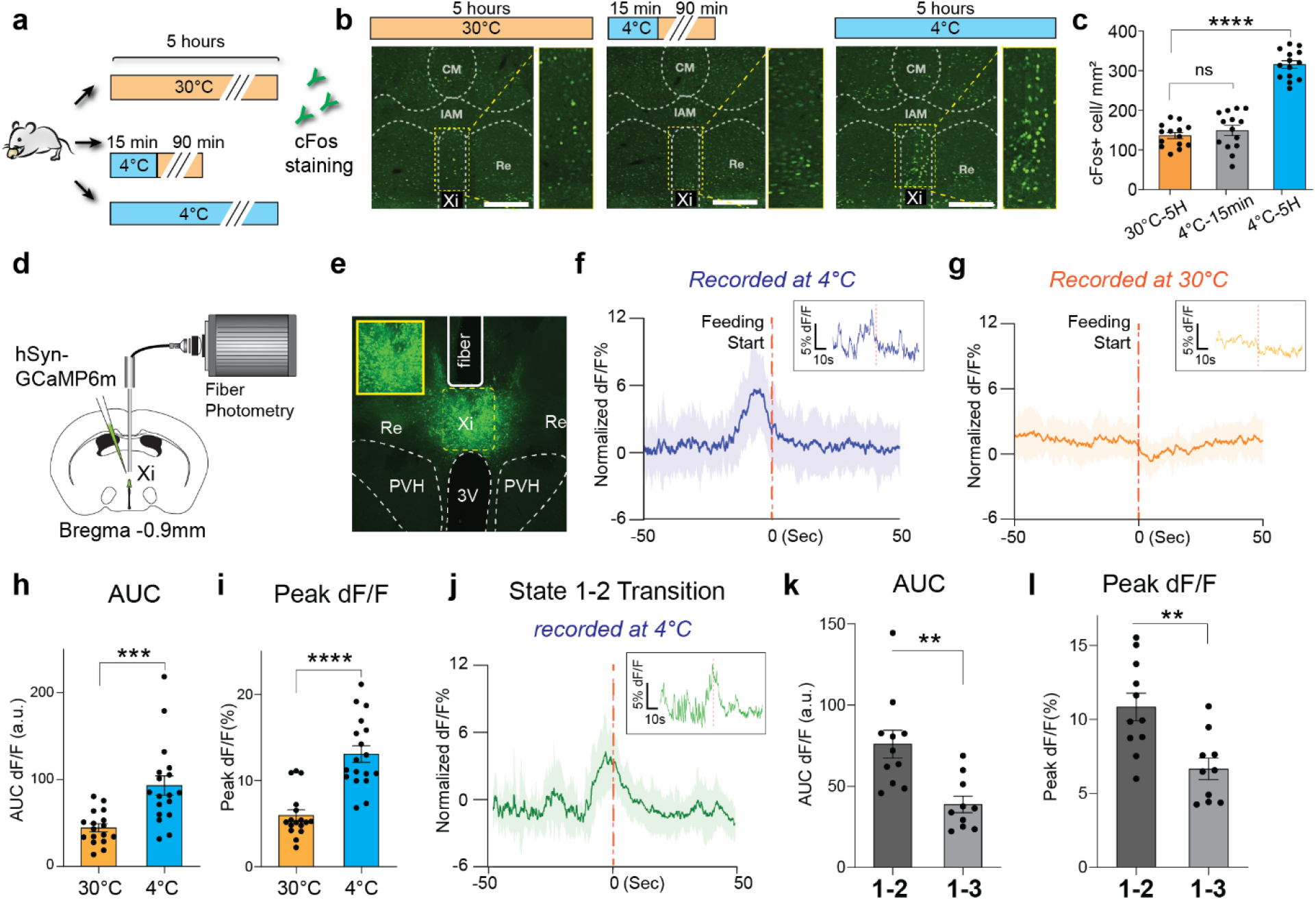
Xi activation is associated with CIEC-induced feeding. **a**, Schematic showing different cold-exposure paradigms used to label cFos expression. **b**, cFos-positive neurons (green) in the vMT after each condition in **a**. Yellow boxes show zoomed-in images of the Xi region. Scale bar 200 μm. CM, central medial nucleus; IAM, intrantereomedial nucleus; Re, nucleus of reuniens; Xi, xiphoid nucleus. **c**, Quantification of cFos+ cells under each condition (n=14 sections from 4 mice for each condition). Data are mean ± SEM. ***P <0.001; ns, non-significant. **d**, Schematic of fiber photometry setup for recording from the Xi. **e**, Representative image of Xi neurons labeled with GCaMP6m below an implanted fiber (solid white line indicates the fiber track). Insert shows zoomed-in view of the Xi neurons. PVH, paraventricular nucleus of the hypothalamus; 3V, 3rd ventricle. **f-g** Fiber photometry signal of AAV-GCaMP6m expressing vMT/Xi neurons in cold (**f)** or thermoneutral (**g**) conditions, shown as the average of 18 events from 4 different mice. The start of the feeding event is indicated by the red dotted line. Insets show an example of a single trace. **h-i** Bar graphs of the area under the curve (AUC) dF/F (−20s to 10s) (**h**) and peak dF/F % (i) for fiber photometry data in **f** and **g. j**, Fiber photometry signal aligned to the state 1-2 transition (red dotted line) based on HMM, averaged from 11 events from 3 different mice. Inset shows an example of a single trace. **k-l** Bar graph of the area under the curve (AUC, -10s to 10s) (**k**) and peak dF/F (**l**), quantified from **j** and Extended Fig. 4. Data are mean ± SEM. ***P =0.004 for AUC in **h**, ****P <0. 0001 for peak dF/F in **i**, **P =0.0017 for AUC in **k**, and **P =0.0024 for peak dF/F in **l** using an unpaired t-test.

### Xi activity is associated with CIEC-induced feeding

To determine how Xi activity represents different behavioral components induced by the cold in real-time, we used fiber photometry to record *in vivo* calcium dynamics of the Xi (Fig. 2d-e). Consistent with the cFos results (Fig. 2b-c), we found that an acute decrease in temperature (23°C to 4°C) did not lead to activation of the Xi (Extended Data Fig. 4e). Next, we analyzed Xi calcium dynamics during CIEC-associated food seeking. Freely moving mice were placed in the cold with food and water for 5 hours before fiber photometry recording. There was a marked increase in Xi activity prior to each feeding event in the cold, but this was not observed in feeding bouts at thermoneutral (non-CIEC) in the same individuals (Fig. 2f-i), suggesting that the Xi activity was selectively associated with CIEC-induced feeding. To gain additional insight into cold-induced Xi activity, we created a scenario with exacerbated cold-induced energy deficit (CIEC+) by briefly restricting food access during the first 3 hours of cold exposure. Under CIEC+ conditions, we observed further elevated Xi calcium transients within the same animals, suggesting Xi activity was scalable with CIEC (Extended Data Fig. 4f-h). Furthermore, Xi neurons did not respond to canonical, fasting-induced feeding either at room temperature or 30°C (Extended Data Fig 5a-c), suggesting their activity is specifically associated with CIEC but not a general energy deficit associated with food restriction. Plasma levels of glucose and leptin, which normally decrease with fasting, were not changed during CIEC (Extended Data Fig 1j, 6a-b); whereas there were elevated levels of lipolysis products (Extended Data Fig. 6c-f), suggesting that noncanonical pathways are signaling the CIEC state to the brain, although the exact molecular nature of such a pathway remains to be determined.

By further incorporating the HMM states into fiber photometry analysis, we discovered that Xi activity is strongly associated with the state transition between the energy-conserving state (state 1) and the food-seeking state (state 2), but not with the transition between state 1 and the non-food seeking, exploratory state 3 (Fig. 2j-l and Extended Data Fig. 4i), suggesting that Xi activity is not associated with general exploratory movements. Together these results indicate that Xi neurons are specifically activated during the transition between states 1 and 2 and leads to the compensatory feeding in a manner that scales with increasing amplitude at higher CIEC.

### Chemogenetic modulation of Xi bi-directionally regulates CIEC-induced feeding

The temporal relationship between Xi activity and CIEC-induced food seeking led us to ask whether Xi neurons could causally modulate this behavior. However, the Xi is a small, less-studied midline thalamic nucleus without well-defined boundaries or molecular markers^23,24^. Noting that CIEC selectively induced cFos in the Xi without activating the surrounding areas (Fig. 2b), we hypothesized that we could target these Xi^CIEC^ neurons based on their unique activation history during prior cold exposures. Leveraging a previously-validated vCAPTURE strategy^17,25–28^ based on activity-dependent ESARE-ER-Cre-ER^29^, we targeted cold-activated Xi neurons with an hM3Dq (Gq -coupled human muscarinic M3 designer receptor exclusively activated by a designer drug (DREADD) or an RFP control (Fig. 3a-c). Consistent with the specificity and efficiency of previous CAPTURE and TRAP work^25,27,29^, we found that Xi^CIEC^ neurons were efficiently captured at 4°C and that these neurons highly overlapped with cold-induced cFos+ neurons (89.9±6.7% by cFos/CAPTURE, 52.6±12.4% by CAPTURE/cFos), whereas minimal neurons were captured at 30°C (Fig 3a, d-f, also see Fig. 2 a-b).

**Fig. 3.**
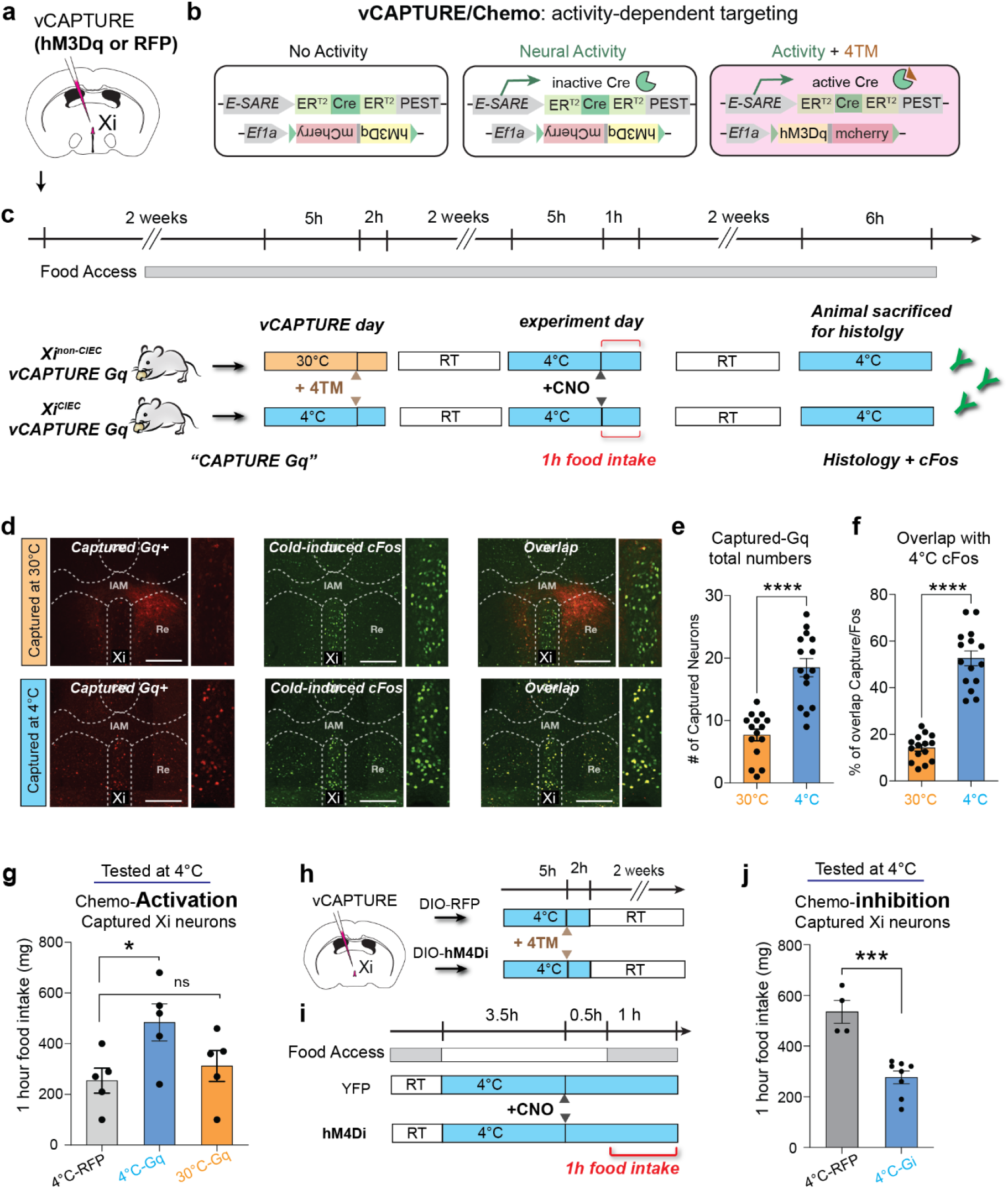
Xi neurons modulate CIEC-induced feeding. **a**, Schematic showing injection site for AAV-eSARE-ER-Cre-ER and AAV-DIO-hM3Dq or RFP control in the Xi. **b**, Schematics of vCAPTURE activity-dependent labeling strategy to express hM3Dq DREADD in CIEC activated Xi neurons. **c**, Schematics showing procedures for activity-dependent capture of the CIEC vs non-CIEC neurons in the Xi. For vCAPTURE of Xi^CIEC^ neurons, Tamoxifen was given to mice 5 hours after onset of cold (4°C) exposure. **d**, Representative histology from non-CIEC (30°C, top) and CIEC (4°C, bottom) mice used for cFos double labeling. Left: vCAPTURE-mediated hM3Dq labeling in the vMT/Xi region; middle: CIEC-activated cFos labeling; right: overlay of vCAPTURE and cFos double labeling. Scale bar: 500 μm **e-f**, Quantification of cFos double-labeling with vCAPTURE neurons in the Xi from **d**. Data are mean ± SEM. ****P <0.0001 using an unpaired t-test. **G**, Bar graph showing food intake levels after activation of Xi^CIEC^ vs. Xi^non-CIEC^ neurons compared to RFP-controls (n=5 mice per group). Data are mean ± SEM. *P =0.0397 and ns=0.7374 using an ordinary one-way ANOVA with Dunnett correction for multiple comparisons. **H-I** Schematics of vCAPTURE strategy and procedure to test loss of function experiment with DREADD-Gi(hM4Di). Note the Gi mice were briefly food-restricted (indicated by gray bars) before clozapine N-oxide (CNO) injection to elevate baseline feeding. **j** Bar graph showing food intake levels after inhibiting Xi^CIEC^ neurons (n=4 mice for RFP and n=4 for Gi). Data are mean ± SEM. ***P =0.0003 using an unpaired t-test.

Upon reactivation by CNO, we observed significantly increased food intake in mice targeted with hM3Dq in the cold (Xi^CIEC^) compared to mice targeted with the RFP control or at 30°C (Xi^non-CIEC^) (Fig. 3g). We then used the same vCAPTURE strategy to target the Xi^CIEC^ with an inhibitory hM4Di DREADD. A significant decrease in CIEC-induced food intake was observed upon CNO-mediated inhibition compared to the RFP-control (Fig. 3h-j, Extended Data Fig. 7a). By contrast, neither chemogenetic inhibition nor activation affected food intake at room temperature (Extended Data Fig. 7b-c), further indicating that the role of Xi neurons is specific to cold-induced feeding.

### Xi^CIEC^ neurons encode an internal state-dependent, context-specific valence

To understand how Xi^CIEC^ neurons promote feeding with higher temporal resolution, we turned to optogenetics. We first injected ChR2-expressing AAV directly into the vMT region and analyzed feeding behaviors in a CIEC scenario (Extended Data Fig. 8a-b). Optogenetics stimulation of the bulk vMT region led to higher food intake but also caused a rapid increase in overall movement that was not observed during the natural cold adaption (Extended Data Fig. 8c-f). These results suggested that non-specific activation of neighboring neurons outside the Xi may have confounded the behavioral output.

To improve the specificity, we again used the activity-dependent vCAPTURE system to selectively express ChR2 in Xi^CIEC^ neurons (Fig. 4a-b). Photostimulation of ChR2-expressing Xi^CIEC^ neurons resulted in a robust increase in food intake without changes in general movement (Fig. 4c-f) or in an open-field test (Extended Data Fig. 9a-d) compared to the RFP controls. Interestingly, in the absence of cold (i.e., at rodent thermoneutrality, 30°C), photostimulation of Xi^CIEC^ neurons only resulted in a minor but non-significant change in food intake (Extended Data Fig. 9e-h), indicating that the Xi^CIEC^ neurons require a CIEC state to induce bona fide feeding in animals. Additionally, video analysis revealed that photoactivation of Xi^CIEC^ resulted in more transitions from the energy-conserving HMM state 1 to the food-seeking state 2, while state 1-3 transitions were not altered (Fig. 4g-h). Similarly, the total time spent in state 2 increased after photostimulation (Fig. 4i, Extended Data Fig. 9i). These results in combination with the earlier fiber photometry data indicate that Xi^CIEC^ activity specifically promotes CIEC-induced feeding without affecting other general movement-related actions.

**Fig. 4.**
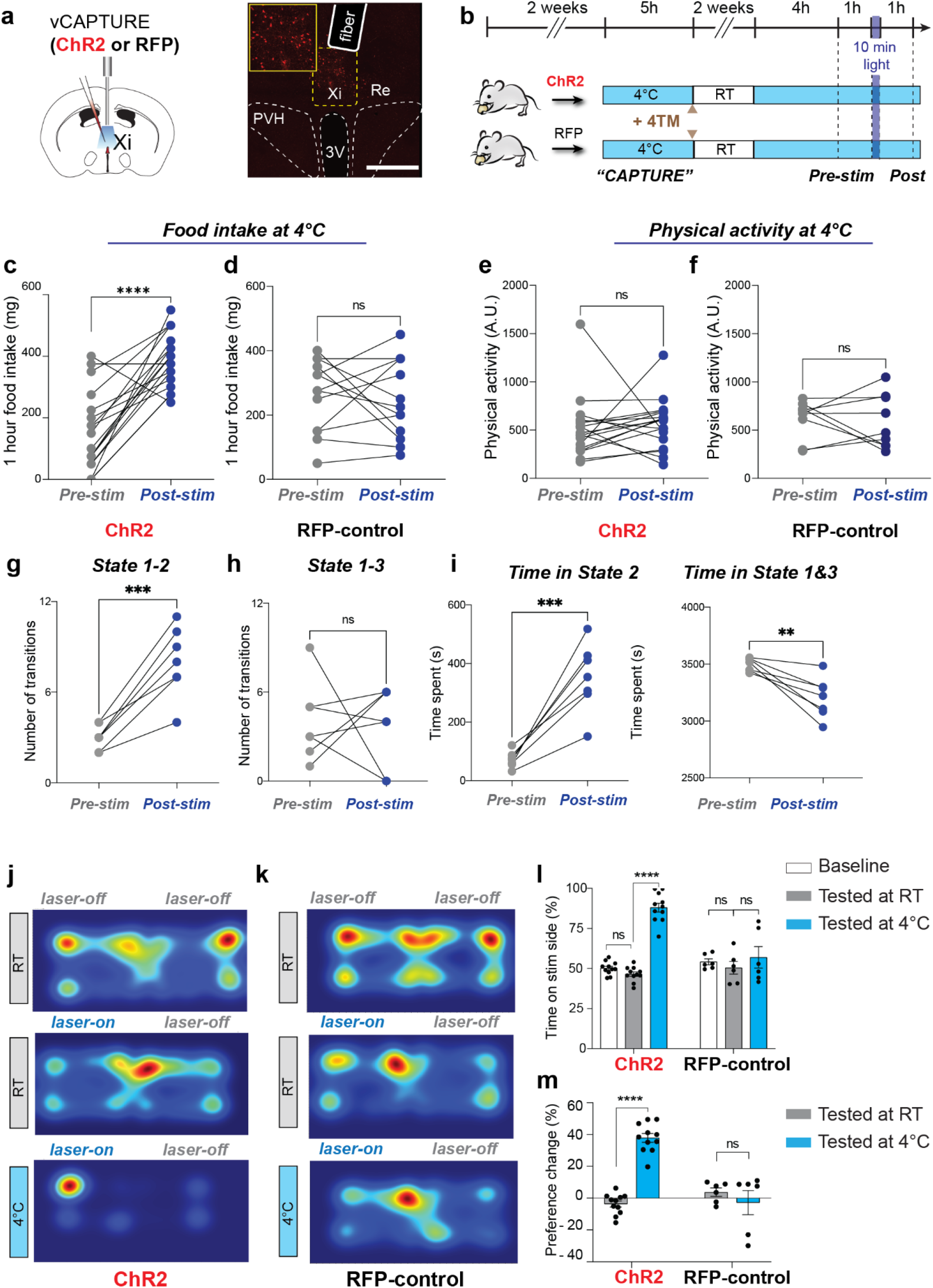
Xi^CIEC^ activity represents a context-specific valence. **a**, Schematic and representative histology showing injection of a viral cocktail consisting of AAV-DIO-ChR2 and AAV-eSARE-ER-Cre-ER, and optogenetic implant at the Xi (solid white line indicates fiber track). Scale bar 500 μm. **b**, Schematics of vCAPTURE optogenetics experimental procedures. **c-d**, Difference in food intake in cold pre- and post-laser stimulation for ChR2 mice (n=9 mice, N=18 times) (**c**) and for RFP control mice (n=7 mice, N=13) (**d**). **e-f**, Difference in physical activity in the cold for ChR2 (n=9 mice, N=18 times) (**e**) and for RFP control mice (n=7 mice, N=9) (**f**). Data are mean ± SEM. ****P < 0.001 for **c**, ns=0.6953 for **d**, ns= 0.6287 for **e**, and ns= 0.7252 for **f** using a paired t-test. **g-h** Total numbers of transitions from state 1 (energy-saving state) to state 2 (exploration with feeding) in ChR2 mice pre- and post-stimulation (n=7 mice) in **g** and from state 1 to state 3 (exploration without feeding) in **h. i**, Total time spent in state 2, pre- and post-stimulation in ChR2 mice (left panel) and total time spent in state 1 and 3 combined, pre- and post-stimulation (right panel) (n=7 mice). Data are mean ± SEM. ***P =0.0005 for **g**, ns =0.8779 for **h**, ***P=0.0006 and **P=0.0046 using paired t-test. **j-m**, Real-time place-preference (RTPP) test based on Xi photostimulation. **j**, Representative heatmap of a ChR2 mouse for basal place preference (no laser, top panel), for laser-on RTPP at room temperature (RT) (middle), and for laser-on RTPP in the cold (bottom). **k**, Representative heatmaps from RFP-controls under the same RTPP conditions. **l**, Bar graph showing quantification of percentage of time spent on the stimulated side. Data are mean ± SEM. ****P <0.0001 for ChR2 base vs ChR2 4°C and ChR2 RT vs ChR2 4°C, ns=0.4589 for ChR2 base vs ChR2 RT, ns=0.7083, 0.9985, or 0.9648 for RFP base vs RT, 4°C or RFP RT vs 4°C respectively, using ordinary two-way ANOVA with Tukey’s multiple comparison test. **m**, Percent change in preference between time spent in stimulation chamber, normalized to no-laser basal level. Data are mean ± SEM. ****P <0.0001 for ChR2 RT vs ChR2 4°C, ns=0.6887 for RFP RT vs RFP 4°C, using ordinary two-way ANOVA with Tukey’s multiple comparison test.

Next, to delineate how Xi neurons affect behavioral transitions, we used real-time place-preference (RTPP) to determine the valence associated with Xi activation. Xi^CIEC^ ChR2-expressing mice were first placed in a two-chamber arena without laser stimulation to get a basal place preference. During the RTPP, Xi^CIEC^ neurons were stimulated when the mice entered one side of the arena and stopped when they crossed to the other side (Fig. 4j-k). At room temperature, Xi^CIEC^ activation did not change place preference. However, when RTPP was conducted in the cold (4°C), the same animals switched their preference to the light-stimulated side (Fig. 4j-m). This result is consistent with our state-dependent activity pattern in earlier fiber photometry studies (Fig. 2f-g). Moreover, such a state-dependent valence switch was only present in ChR2-expressing mice but not in RFP-controls. The RTPP results suggest that Xi-mediated state transition for food seeking during CIEC could be driven by a positive valence.

### Xi to NAc output mediates CIEC-associated food seeking

Lastly, we sought to characterize the cell types and projection targets of the Xi. First, by using vGLUT2-Cre and vGAT-Cre mice^30,31^, we found that Xi-regulated food seeking was primarily mediated by glutamatergic neurons in the Xi but not through GABAergic cells (Extended data Fig 10 a-l). Next, we injected mCherry-expressing AAV to map the projection targets of Xi. Consistent with previous reports^23,24^, Xi projected to the nucleus accumbens (NAc), basolateral amygdala (BLA), and anterior cingulate cortex (ACC) (Fig. 5 a and b). To determine if one or more of these projections corresponded to the Xi^CIEC^ population, we designed a double-labeling experiment in which retrograde CTB dyes were individually injected into the NAc, BLA, and ACC. We found that the highest overlap between cold-activated cFOS and CTB was in the Xi-NAc projecting neurons (Fig. 5d-g), prompting us to test the role of Xi-NAc projection in cold-induced feeding. We injected ChR2-expressing AAV into the Xi and implanted an optic fiber above either the NAc, BLA, or ACC (Fig. 5i, j top panel and Extended Data Fig. 10 a). Photoactivation of Xi-NAc projection, but not Xi to ACC or BLA projection, resulted in a significant increase in food intake (Fig. 5i-j and Extended Data Fig. 11a and b). These projection-specific effects were further confirmed by using two fiber implants (NAc and BLA) in the same animals but photostimulated sequentially (Extended Data Fig. 11c-g). Finally, by RTPP, we also found that activation of Xi to NAc projection resulted in a cold-dependent positive valence, whereas the activation of other projections failed to do so (Fig. 5 k and l). Together these results show that Xi to NAc projection primarily mediates cold-induced food-seeking behaviors.

**Fig. 5.**
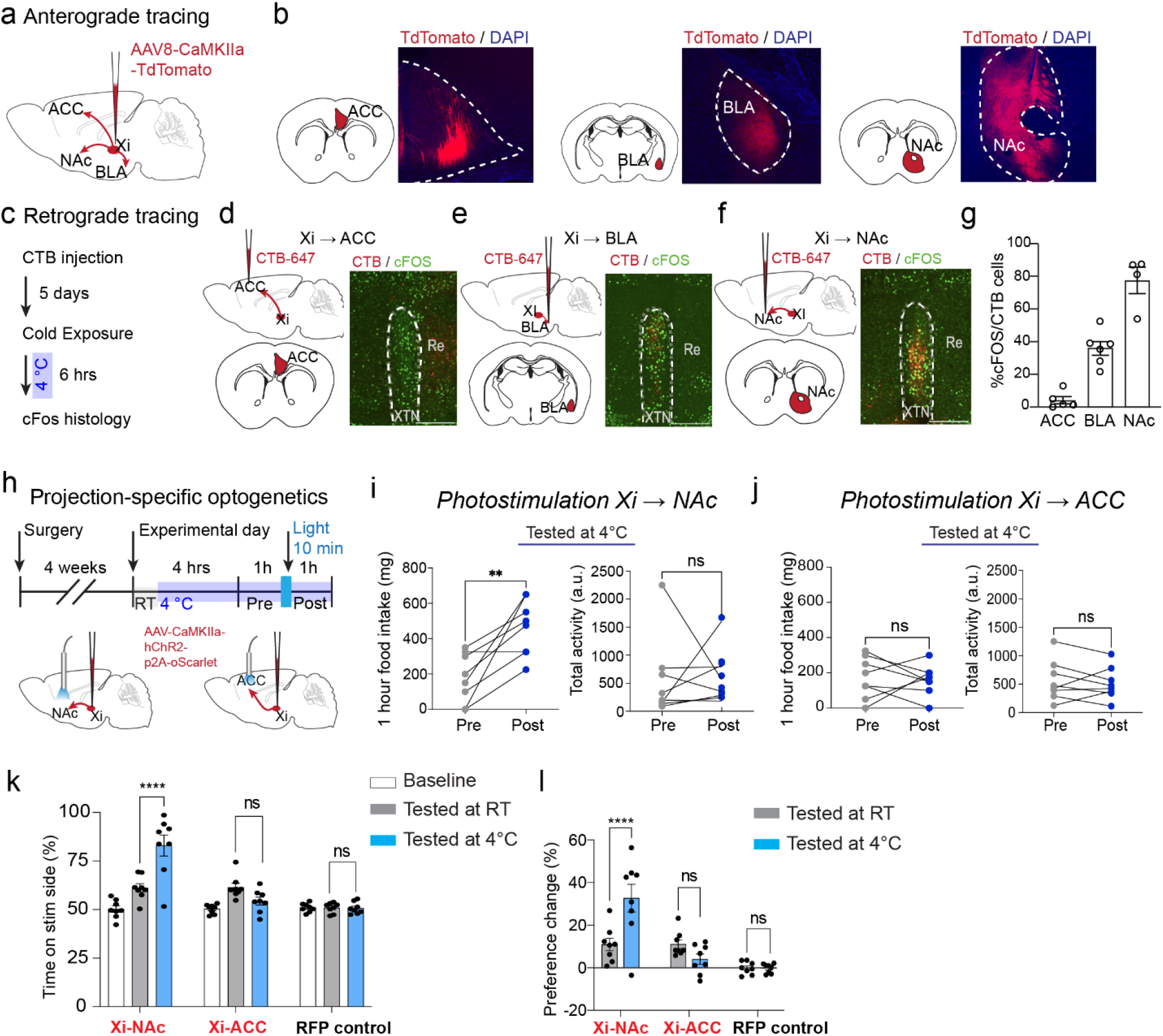
Xi to NAc projection mediates CIEC-associated food seeking. **a-b**, Schematic for anterograde Td tomato labeling. Mice were injected with AAV-CaMKIIa-TdTomato at the Xi, and axonal fibers were observed in the Anterior cingulate cortex (ACC), Basolateral amygdala (BLA), and Nucleus accumbens (NAc) (**b**) (n=5 mice). **c**, Schematic of retrograde CTB and cFOS double labeling experiment. **d-f**, Schematics and representative overlay images between retrograde CTB and cFOS signals at the Xi. **d**, (ACC-Xi), **e**, (BLA-Xi), and **f**, (NAc-Xi). **g**, Quantification of the CTB/cFOS overlap. **h**. Schematic of projection-specific optogenetics experiments. Quantification of change in food intake and physical activity upon stimulation of Xi-NAc (**i**) or Xi-ACC (**j**) projection during CIEC (n=8 mice). Data are mean ± SEM. **P <0.0052 for food intake, ns=0.9163 for physical activity for Xi-NAc in (**i**), ns=0.8037 for food intake and ns=0.5893 for physical activity for Xi-ACC in (**j**) using a paired t-test. **k-l**, Projection-specific RTPP test. RFP control was injected in the Xi and stimulated at the Xi to control for all the projections. **k**, Bar graph showing percentage of time spent on laser stimulation side. Data are mean ± SEM. ****P < 0.0001 for Xi-NAc RT vs Xi-NAc 4°C, ns=0.3027 for Xi-ACC RT vs Xi-ACC 4°C, ns>0.99 for RFP RT vs RFP 4°C. **l**, Percent change in preference upon stimulation, normalized to baseline place preference for the same animal. Data are mean ± SEM. ***P <0.0003 for Xi-NAc RT vs Xi-NAc 4°C, ns=0.6413 for Xi-ACC vs Xi-ACC 4°C, and ns>0.99 for RFP RT vs RFP 4°C, by ordinary two-way ANOVA with Tukey’s multiple comparison test.

## Discussion

Here, leveraging whole-brain screening, in vivo calcium imaging, and chemo- and optogenetic manipulations, we demonstrated that the Xi nucleus serves as a key brain region for promoting cold-induced food-seeking behaviors. Although cold-induced feeding has been widely reported across animal species, including humans, our study suggested that such a feeding increase was not a direct response to cold sensation or a unilateral increase of food seeking behaviors, but rather, a dynamic outcome of the transitional priorities between energy conservation and replenishment in response to elevated caloric debt in the cold.

Interestingly, Xi activity was associated with behavioral transitions (Fig. 2j), but it returned to baseline before the feeding began (Fig. 2f), suggesting that sustained Xi activity is not required for feeding. Recently, the vMT/Xi has been shown to gate behavioral shifts between saliency-reducing and saliency-enhancing strategies in response to visual threats in mice^25^. Reminiscent of this shift, we speculate that the endogenous Xi activity might gate the behavioral transitions from energy conservation to a replenishing (feeding) state, as experimental activation of Xi increased CIEC-associated feeding (Fig. 3g, Fig. 4c), potentially by lowering the threshold to increase the probability of state 1-2 transitions (Fig. 4g-i). This gating, however, is specific to the cold context in which coordinating energy expenditure and intake is critical for survival; it appears to be less involved in starvation conditions, in which the drive for feeding is more unilateral (Extended Data Fig 5). Together, we propose that the Xi could represent an important mechanism for dynamically switching opposing survival strategies during specific natural behaviors.

Feeding behaviors are tightly regulated by both homeostatic and reward circuits^32–34^. The Xi is a less-studied brain region that is not a direct part of the classic hypothalamic or limbic reward systems. However, it projects to multiple well-known regions involved in feeding regulation, including the prefrontal cortex, BLA, and NAc^21^ (Fig 5a-b). We found that Xi-regulated feeding and valence switching behaviors were primarily mediated by the Xi to NAc projection. As a limbic reward center, the NAc is known to integrate motivational and sensory signals to guide motor output^35,36^ and regulate feeding^37,38^. Based on our findings, we propose that the NAc is a key downstream target of the Xi in motivating animals to switch behavioral states as dictated by the internal metabolic state. It remains to be determined, however, how Xi activity interacts or integrates with other aspects of the canonical feeding framework and, especially, what the identity of its input signals are.

Although feeding is one of the most studied rodent behaviors, it has been primarily modeled in the context of food restriction (e.g., overnight fasting) for creating a negative energy balance. Cold-induced feeding provides a new paradigm to study appetite that is driven by energy expenditure. Although both food restriction and energy expenditure create a net negative energy balance, the organismal physiology can be sharply different—for example, fasting is associated with lower blood glucose, leptin, insulin, and suppressed thermogenesis and basic metabolic rates, whereas cold adaptation is accompanied by high glucose utilization, elevated thermogenesis, and higher metabolic rates. Cold-induced energy compensation and associated neural substrates (such as the Xi and many others revealed by the whole-brain screen) could provide an entry point to understand naturalistic energy deficit created by other high energy expenditure states such as exercise and lactation, findings which will be important not only for understanding fundamental mammalian biology but also for obesity and weight management.

## Acknowledgments

We thank J Read, A Brashears, and A Kanow for technical assistance, H Wang and H Zhu for initial behavioral setup, Y Wang for illustration, Scripps Research Department of Animal Resources for animal resources, and members of Ye lab for discussion. We thank L Stowers, C Kim, V Augustine, and W Hong for feedback on the manuscript. We are grateful to AV Kravitz for FED3 applications. NKL, TQ, ZP, and DY were supported by the Dorris Scholar Award. NKL was supported by the AHA postdoctoral fellowship and Dorris Scholar Award. LY is supported by National Institutes of Health Director’s New Innovator Award (DP2DK128800), NIDDK (DK114165, DK124731), the Dana Foundation, the Whitehall Foundation, Baxter Foundation, and the Abide-Vividion Endowment. ASB is supported by DK133948, DK107717, and OD028635.

## Author contributions

Conceptualization: NKL, LY. Investigation and Analysis: NKL, PL, SA, AZ, KW, TQ, ZP, DY, VN, ASB. Funding acquisition: LY. Project administration: LY. Supervision: LY, GWY, ASB. Writing – original draft: NKL, LY, Writing – review & editing: All authors.

## Data and materials availability

All data are available in the main text or supplementary materials.

## Methods

### Mice

Animal experiments were performed either at Scripps Research Institute, La Jolla or Beth Israel Deaconess Medical Center (BIDMC). Experiments were approved by the Scripps Research Institute’s or BIDMC’s Institutional Animal Care and Use Committee (IACUC), respectively. Wild-type C57BL/J6 mice were injected with viruses between 6-8 weeks of age and were allowed to recover for at least one week before experiments or experiment-related habituation. Group size was not pre-determined based on statistical power law but on previously published studies. Animals were randomly assigned to different experimental groups before viral injections. Both male and female were used for histology and cFos labeling. Male mice were used for free-moving behavioral studies.

### Reagents and Viruses

AAV8-hSyn-DIO-Gq–mCherry (Addgene, 44361-AAV8), AAV8-hSyn-DIO-Gi–mCherry (Addgene, 44362-AAV8), AAV8-hSyn-mCherry (Addgene, 114472-AAV8), and AAV9-Syn-GCaMP6m-WPRE-SV40 (Addgene, 100841-AAV9) were obtained from Addgene.

AAV5-ESARE-ER-Cre-ER-PEST was obtained from UNC, AAV8-Ef1a-DIO hChR2(H134R)-p2AScarlett and AAV8-CaMKIIa-hChR2-p2A-oScarlet were obtained from Stanford. cFos antibody was purchased commercially (Cell signaling Cat # 2250) as was CNO (Hello bio Cat # HB1807).

### Stereotactic viral injection and fiber implantation

Mice were anesthetized with 3.0% isoflurane in an induction chamber, and after 5 minutes, the animals’ heads were fixed on a stereotactic device (Kopf) using ear bars. Mice were maintained under 0.8-1.2% isoflurane during surgery on a warm heating pad (body temperature was maintained at 36°C). The heads were shaved to remove hair, and using a scalpel blade, a midline incision was made to expose the skull. The mouse head was balanced using a bregma and lambda system on the stereotaxic headstand. Using dental drills, small craniotomies were made above the site of injections. The virus was infused at 75 nl min−1 using a Nanoject II (Drummond) injector connected to a 10-μl Hamilton syringe. The needle was kept at the injection site for 10 minutes before withdrawing. Mice were allowed to recover for one-two weeks before experiments or experiment-related habituation.

For chemogenetic stimulation, a cocktail of 125 to 150 nl of AAV8-hSyn-DIO-Gq–mCherry or AAV8-hSyn-mCherry and AAV5-ESARE-ER-Cre-ER-PEST was delivered to the vMT/Xi region (bregma: −1.0 mm, midline: 0.0 mm, dorsal surface: −4.4 mm). The final concentration of ESARE viruses in the cocktail was 7×10^12^ GC per ml. Two weeks after surgery, mice were habituated to TM injection (explained below in the vCAPTURE protocol section).

For optogenetics experiments with non-selective activation of the whole vMT/Xi region (bregma: −1.0 mm, midline: 0.0 mm, dorsal surface: −4.4 mm), 150 nl of AAV8-CaMKIIa-hChR2-p2A-oScarlet (5×10^12^ GC per ml) was injected at the vMT/Xi region. After removing the needle, an optical fiber cannula (200-μm diameter, 0.22 N.A., RWD life sciences) was implanted at the vMT/XI, 100-200 μm dorsal to the injection site at (bregma: −1.0 mm, midline: 0.0 mm, dorsal surface: −4.2 to -4.3 mm). The optical cannula was cemented to the skull with dental cement. For the activity-dependent optogenetics labeling cohort, 200 nl of a cocktail of AAV8-Ef1a-DIO hChR2(H134R)-p2AScarlett + AAV5-ESARE-ER-Cre-ER-PEST (7×1012 GC per ml) was injected at the vMT/Xi region.

For fiber photometry experiments, 150-180 nl of AAV9-Syn-GCaMP6m-WPRE-SV40 (7×10^12^ GC per ml) virus was injected using stereotaxic surgery at the vMT/Xi region (bregma: −1.0 mm, midline: 0.0 mm, dorsal surface: −4.4 mm) and an optical fiber cannula (400-μm diameter, 0.5 N.A., RWD life sciences) was implanted at (bregma: −0.9 mm, midline: 0.0 mm, dorsal surface: −4.2 mm). For both optogenetics and fiber photometry, only a single fiber optic cannula was implanted because the Xi is a midline structure.

### Cold and warm exposure

A temperature-controlled rodent incubator maintained at 4°C (RIS28SSD, Powers Scientific) was used for cold exposure. For thermoneutrality, a temperature-controlled warm room set to 30°C was used. For all temperature exposure experiments, all mice were individually housed with a small amount of bedding materials to ensure each individual received the same temperature exposure. For cold exposure, animals were directly transferred into the 4°C incubator in a lidless cage to facilitate the temperature equilibrium. Mice were provided free access to chow food and water during most cold or warm exposure experiments, except in the CIEC+ condition and during the inhibitory chemogenetic experiments, in which they were food-restricted for various amounts of time as indicated in each figure.

### Optogenetic, chemogenetic, and food intake measurement

Mice were allowed to recover for 7 days after optogenetic implant surgery for the non-selective optogenetics experiment or 14 days after tamoxifen (TM) injection for the vCAPTURE cohort of mice. Optogenetics mice were trained and habituated to pellets (20 mg pellet, cat#F0071, Bioserv) along with dispensing systems FED2 (1) in their home cage for 5 days before giving free access to food pellets for 4 days to avoid developing a hedonic value for food pellets. The mice were further habituated to the behavioral chambers for 5 days with food pellets and pellet dispensing systems FED2 before the first experiment. All food intake experiments were performed between 8:00 am to 3:00 pm to avoid the natural circadian cycle influence on feeding behavior.

For the optogenetics experiments, the experimental mice were connected to a blue light laser (473 nm) with an optical fiber (200-μm diameter, 0.22 N.A.; RWD). The light was delivered in 10 ms pulses at 20Hz, 3 sec on, 2 sec off, for 10 minutes. The light power at the fiber tip was 10 mW. For chemogenetics experiments, a regular chow pellet was used for measuring food intake by weight to increase the throughput. The stock solution of CNO was prepared freshly by dissolving in DMSO. Then, the stock solution was diluted in saline, and mice were injected with 3 mg/kg CNO for activating the Gq cohort and 5 mg/kg for the inhibitory Gi cohort. Control mice were given the same amount of CNO as the experimental cohort.

### Indirect calorimetry

Metabolic cage data on 24 individually housed mice were placed in a Promethion indirect calorimeter (Sable Systems) with a temperature-controlled cabinet (Pol-Eko) and provided with ad libitum food (Labdiet 5008, 3.56 kcal/g) and water purified by reverse osmosis. Mice were maintained under a 12-hour/12-hour light/dark photoperiods (0600-1800) at an ambient temperature of 23 ±0.2°C. Temperature transitions from 23 to 4°C were performed over 3 hours starting at 0600. Data were exported with Macro Interpreter, macro 13 (Sable Systems) prior to analysis in CalR version 1.3 (2). Rates of energy expenditure were calculated with the Weir equation (3). Two datasets were analyzed. Dataset 1, 24 mice were maintained at 23°C for 2.5 days (62 hr). The 4°C ambient temperature was maintained for an additional 9 hours before returning to 23°C for 16 hours. Dataset 2: 11 mice were maintained at 23°C for 5 days (115 hr) followed by 4°C for another 5 days (118 hr), Extended Data Figure 1c-e.

### vCAPTURE labeling of Xi CIEC-associated neurons

To label CIEC-associated neurons in the vMT/XI region with activating Gq-DREADD, the mice received a stereotaxic injection of viral cocktail with ESARE-ER-Cre-ER + DIO-Gq and were habituated with the setup every day for 2-3 hours for one week. The ESARE-ER-Cre-ER /Gq mice were randomly divided into XiCIEC and Xinon-CIEC groups. On the day of labeling, the XiCIEC group of mice were placed in the cold for 6 hours with ad libitum food and water. Mice were given 20 mg/kg 4-hydroxytamoxifen (4-TM) (Sigma H6278) through intraperitoneal (i.p.) injection. Mice were injected 6 hours after being in the cold. When these mice enter CIEC, the vMT/Xi neurons that are active during CIEC induce FOS and CreERT2. After 4-TM i.p injection, the CreERT2 will selectively recombine the AAV8-hSyn-DIO-Gq–mCherry encoding Cre-dependent Gq-DREADD in the cell that was expressing FOS. This recombination leads to permanent expression of Gq-DREADD in those cells, which can then be selectively activated at a later point by administration of DREADD agonist clozapine N-oxide (CNO). After the 4-TM injections, these mice were kept in the cold for another 2 hours before returning them back to their home cage kept at 23°C. As a control, we generated another cohort of mice that received a stereotaxic injection of the same viral cocktail as XiCIEC and were habituated using the same protocol, except that on the day of labeling, they received the 20 mg/kg of 4-TM injection after being in 30°C temperature for 6 hours rather than in the cold. cFOS data in Figure 1 showed that very few cells were active in the vMT/Xi region while mice were in thermoneutral conditions for 6 hours compared to being in the cold. We call this cohort of mice “ Xinon-CIEC”. Both sets of mice were injected at the same time of day to avoid labeling variability associated with circadian rhythm. A similar protocol was followed for the optogenetics mice that received a stereotaxic injection of a viral cocktail of AAV5-ESARE-ER-Cre-ER-PEST and AAV8-Ef1a-DIO-hChR2(H134R)-p2AScarlett.

### Histology

Mice were anesthetized with isoflurane before transcardial perfusion with ice-cold PBS followed by cold 4% paraformaldehyde (PFA). Brains were removed and post-fixed overnight with 4% PFA at 4°C. The next day fixed brains were submerged in 3% agarose and kept in cold for 2 hours for embedding. Brains were sliced on a vibratome (Leica VT1000S) into 80-μm coronal sections and collected as two equal sets in a 6-well plate filled with PBS. For IHC staining permeabilization, brain sections were incubated with a blocking buffer containing PBS, 0.3% TritonX-100 (PBST), and 5% donkey serum for 1 hour at room temperature with gentle shaking. After permeabilization, brain sections were incubated with primary antibodies at 4°C in a blocking buffer for 16 hours. Next, brain slices were washed three times with PBST for 20 minutes each to wash away the unbound primary antibody and incubated with secondary antibodies for 1 hour at room temperature diluted in blocking buffer. Finally, brain slices were washed again with PBST 3 times, 20 minutes each, to wash away the unbound secondary before mounting onto super-frost Plus glass slides (VWR). Brain slices were imaged using Olympus FV3000 confocal microscope with 10X objective, 0.6 NA, water immersion (XLUMPlanFI, Olympus). Each brain slice was imaged at 10 μm Z stack. cFOS quantification was done for a single plain (numbers/mm2). For double labeling cFOS vCAPTURE quantification, cells were counted in a region of interest (ROI) defined manually following anatomical landmarks.

### Whole-brain imaging and analysis

Wild-type C57BL/J6 mice were habituated to the behavioral setup for 5 days before the actual experiment. On the day of the experiment, individually housed mice were kept in either cold 4°C or thermoneutral 30°C conditions for 6 hours with free access to food and water before perfusion. Brains were harvested and fixed overnight in 4% PFA. Whole-brain clearing, imaging, and automated analysis were performed by LifeCanvas Technologies through contracted service. Briefly, fixed whole brains were prepared with SHIELD to preserve protein antigenicity before being actively cleared and immunolabeled with a cFos antibody using SmartBatch+. Labeled brains were index-matched by EasyIndex and imaged by volumetric lightsheet microscopy (SmartSPIM), followed by image post-processing, cell quantification, atlas registration, and regional graphics. The averaged numbers of cFOS+ cells in each region were grouped into seven “ buckets” based on ABA annotations for data visualization. The size of the dots represents the mean differences between 30°C and 4°C conditions in each region.

### Fiber photometry

Mice were attached to a patch fiber (400-μm core NA 0.5, RWD) connected to a 470 nm and 410 nm light source. Fiber photometry and acquisition setup has been previously described (2). 470-nm excitation light was used to measure the Ca^2+^ signal, and a 410-nm excitation light was used as a reference signal. Changes in Ca^2+^ fluorescence (470 nm) signal was compared with background Ca^2+^ fluorescence (410 nm), providing an internal control for movement and signal bleaching. sCMOS camera (Hamamatsu, Orca Flash 4.0 v2) was used to capture the patchcord end-face images and processed using previously described MATLAB code (4). Data-acquisition hardware (National Instruments, NI PCIe-6343-X) was used to digitize the images at 5 kHz. Signals were expressed in delta F/F, where F represents baseline fluorescence. Video recording of behavior was used to manually annotate and timestamp feeding and other behaviors. AUC and peak values were calculated for -20/-10 to 10 seconds from the defined event (for feeding/HMM states, respectively).

For fasting-refeeding experiments, mice were fasted for 16 hours. Mice were placed in a tall chamber and their cannula was attached to the fiber photometry setup. Mice were allowed to habituate for 30 minutes and a feeding device (FED3) was placed at one corner of the chamber. FED3 was programmed to start dispersing pellets after 15 minutes of placing the device in the chamber to avoid introducing noise in the experiment. Data acquisition, analysis, and processing were done as described above.

### Measurement of serum leptin, glycerol, and free fatty acid

20 mice were divided into 5 groups: 1 (2 hour Cold), 2 (4 hour Cold), 3 (2 hour Thermoneutral), 4 (4 hour Thermoneutral), and 5 (home cage 0 hour). Quantification of blood plasma measurement was performed according to the manufacturers’ recommendations (Mouse Leptin ELISA kit Crystal Chem (90030), Free fatty acid quantification kit Abcam (ab65341), Free glycerol assay kit abcam (ab65337)).

### Hidden Markov Model

Wild-type C57BL/6 mice were habituated to the setup (using FED2) for 4 days before recording their behaviors at 4°C. For each mouse, two video were recorded, one from the top view and one from the side view using two separate Logitech C270 web cameras. We manually annotated the start and end of each behavior. Analyzers were blind to the experimental groups of mice. The behaviors include sitting, shivering, grooming head, turning, lower body grooming, moving out, eating, moving back, pushing bedding, standing up, bedding retrieval, drinking, digging, grooming tail, and walking. The behaviors were annotated for every second. Hidden Markov Model (HMM) requires an input sequence, an estimated transition matrix, and an estimated emission matrix. The input sequence was generated with data from the 180 minute videos, previously analyzed manually, of mice exposed to cold for 4-6 hours with free access to food and water. The input sequence of each mouse was put into a vector. We defined a 3-state HMM: (1) energy-conserving state, (2) exploration with food consumption, and (3) exploration without food consumption. We generated the initial guess for the estimated transition matrix by assuming equal probabilities for all transitions as we did not have a priori information. We generated the initial guess for the estimated emission matrix by predicting the probability of behavior in each state. We used a custom MATLAB code using the Baum-Welch algorithm to get the transition and emission matrices. To calculate the underlying state of behavior based on a sequence of behaviors, we used the MATLAB built-in function of hmmviterbi.

### Real-time place preference (RTPP)

For basal place preference (BPP) measurements of activity-dependent optogenetics mice, ChR2-XiCIEC or RFP-XiCIEC were placed in a two-chamber acrylic box (60 cm length × 25 cm width × 30 cm height, with each side of the chamber measuring 30 cm × 25 cm) at room temperature (23°C). Each chamber had different contextual pattern cues on the wall. One side had a black and white stripe pattern on the wall, and the other had a dotted pattern. Mouse movements were recorded with an overhead top view Logitech web camera for the entire 30-minute session. Mice were habituated for several hours daily with fiberoptic cable attached to their head implants for a week in a tall chamber before BPP. Videos were manually quantified for the amount of time spent in each chamber. For measuring real-time place preference (RTPP) at room temperature, mice were placed in the center of the place preference chamber, and the laser was switched on when all four paws of the mouse were in the stimulation chamber. The laser was manually switched on (10 mW, 10 ms, 20 Hz) and off by the operator sitting in the next room while monitoring a live overhead video of the mouse. The laser was kept on while they remained in the stimulation chamber and switched off when they left the stimulation chamber. To test the change in valence while mice are undergoing CIEC, mice were placed at 4°C for 5 to 6 hours with access to ad libitum food and water before performing RTPP in the cold. Mice were given a one-week gap between RTPP at 23°C and 4°C.

### Open-field test

To measure anxiety-like behavior in the optogenetics cohort, mice were placed in the outer zone of the white-walled acrylic open-field arena (60 cm × 60 cm), and movement was recorded for 20 minutes using a top view Logitech web camera. The laser was turned on and off intermittently for a 3-minute period by an operator sitting in the next room while monitoring a live video feed. Total time and distance in the center 40% of the area were analyzed using custom automated tracking software. The open field test was performed only once for each mouse. Mice were returned to their home cage after the session.

### Statistics

Two-way ANOVAs were used to assess how behavior was affected by other factors (e.g., RTPP tested by optogenetic manipulations and temperature). Unpaired t-tests were used for comparisons between the two groups. Two-tailed tests were used throughout with α = 0.05. Multiple comparison adjustments, biological replicates, and significance definitions are included in each figure legend.

