## Supplement_Figs for "Xiphoid nucleus of the midline thalamus controls cold-induced food seeking"

**Extended Data Figure 1**

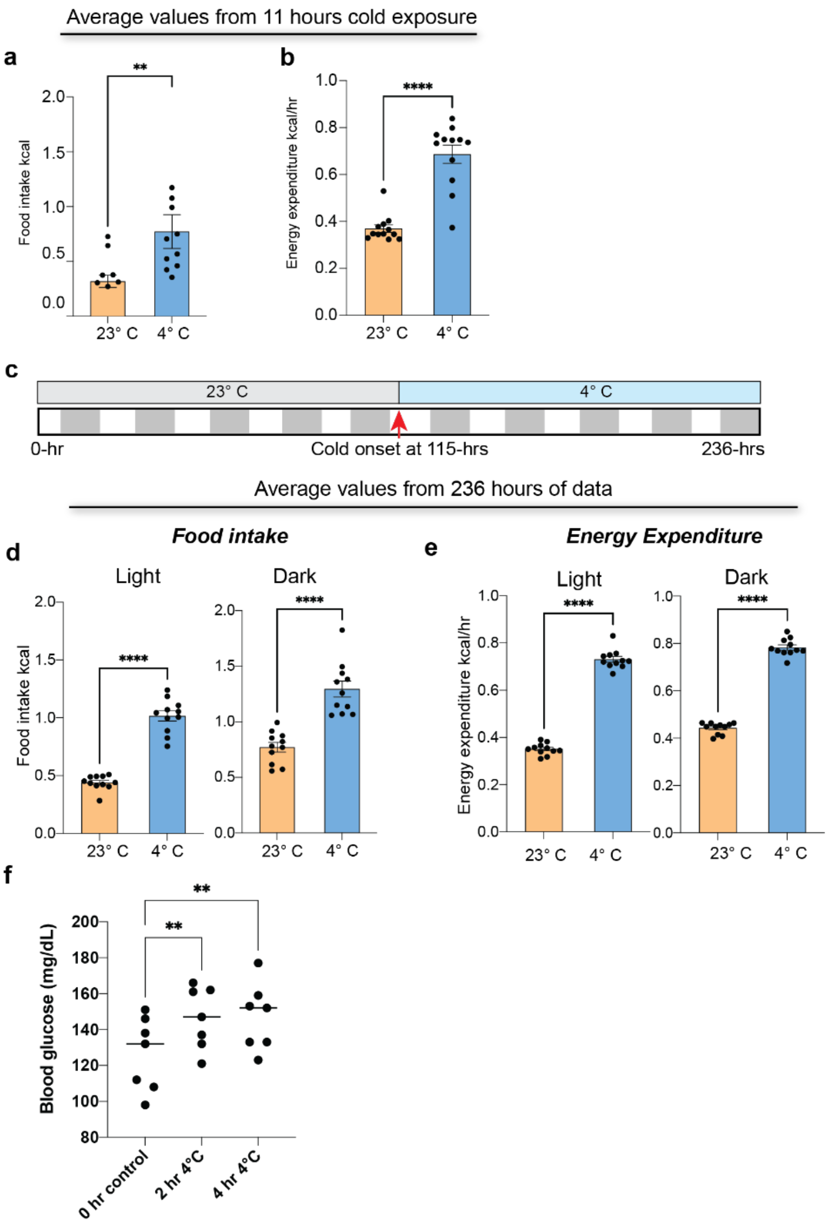


**Extended Data Fig. 1| Metabolic changes during CIEC. a-b,** Bar graphs showing average food intake (**a**) and energy expenditure (**b**) for data presented in Fig 1a-c. c. Schematic depicting the long-term cold exposure protocol**.** 5 days (115 hrs) after being at RT (23°C), the temperature was switched to 4°C. white: light phase, grey: dark phase. **d-e,** Quantification of food intake (**d**) and energy expenditure (**e**) during light and dark phases before and after the temperature switch. For calculating the average food intake and energy expenditure, values from either 4 days or 4 nights were used for both RT and 4°C (excluding the day of temperature switch). Data are mean ± SEM. ****P < 0.0001 using a paired t-test. **f,** Blood glucose levels of WT mice at 2 hrs and 4 hrs after exposure to cold (timepoints are from the same animals). Data are mean ± SEM. **P = 0.001 for comparison between 0 hr and 2 hr 4°C, **P = 0.0028 for 0 hr vs 4 hr 4C, using a one-way ANOVA with Dunnett’s multiple comparison test.

### Extended Data Figure 2

**a**

Estimation of Emission Matrix (Percentage)

| | Sit | Shiver | Groom head | Turn | Lower body groom | Move out | Eat | Move back | Push bedding | Stand up | Bedding retrieval | Drink | Digging | Groom tail | Walk |
|---|---|---|---|---|---|---|---|---|---|---|---|---|---|---|---|
| State 1 (Energy conserving) | 87 | 6 | 2.1 | 0.9 | 1.9 | 0 | 0 | 0.57 | 0.54 | 0 | 0 | 0 | 0 | 0.32 | 0 |
| State 2 (Exploration with food seeking) | 0 | 0 | 0 | 0 | 0 | 24.5 | 74.9 | 0 | 0 | 0.58 | 0 | 0 | 0 | 0 | 0 |
| State 3 (Exploration without food seeking) | 0 | 0 | 0 | 0 | 0 | 3.58 | 0 | 0 | 0 | 19.6 | 23.2 | 13.8 | 16.7 | 0 | 23.2 |

**b**

Estimation of Transtion Matrix (Percentage)

| | State 1 (Energy conserving) | State 2 (Exploration with food seeking) | State 3 (Exploration without food seeking) |
|---|---|---|---|
| State 1 (Energy conserving) | 99.56 | 0.21 | 0.23 |
| State 2 (Exploration with food seeking) | 4.71 | 93.55 | 1.75 |
| State 3 (Exploration without food seeking) | 6.50 | 2.22 | 91.28 |

**c**

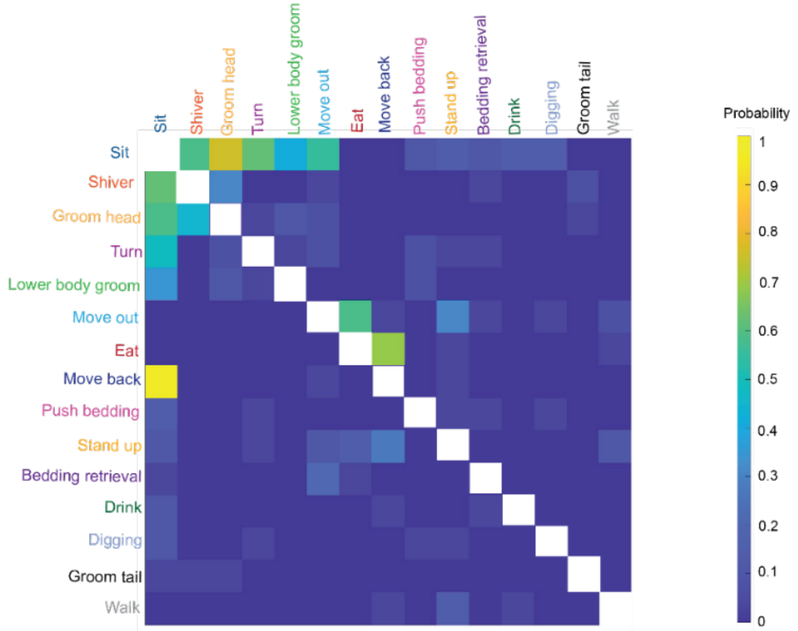


**Extended Data Fig. 2| Representation of Hidden Markov model (HMM) of CIEC paradigm in wild-type mice. a,** Using 15 different labels of actions taken by mice in the CIEC paradigm, a HMM estimated the emission matrix with three states (energy conserving (state 1), exploration with food-seeking and consumption (state 2), and exploration without food-seeking (state 3)). (n=3 mice). **b,** Corresponding transition matrix probabilities between each state. **c,** Transition matrix of different actions taken by the mouse during the CIEC state.

**Extended Data Figure 3**

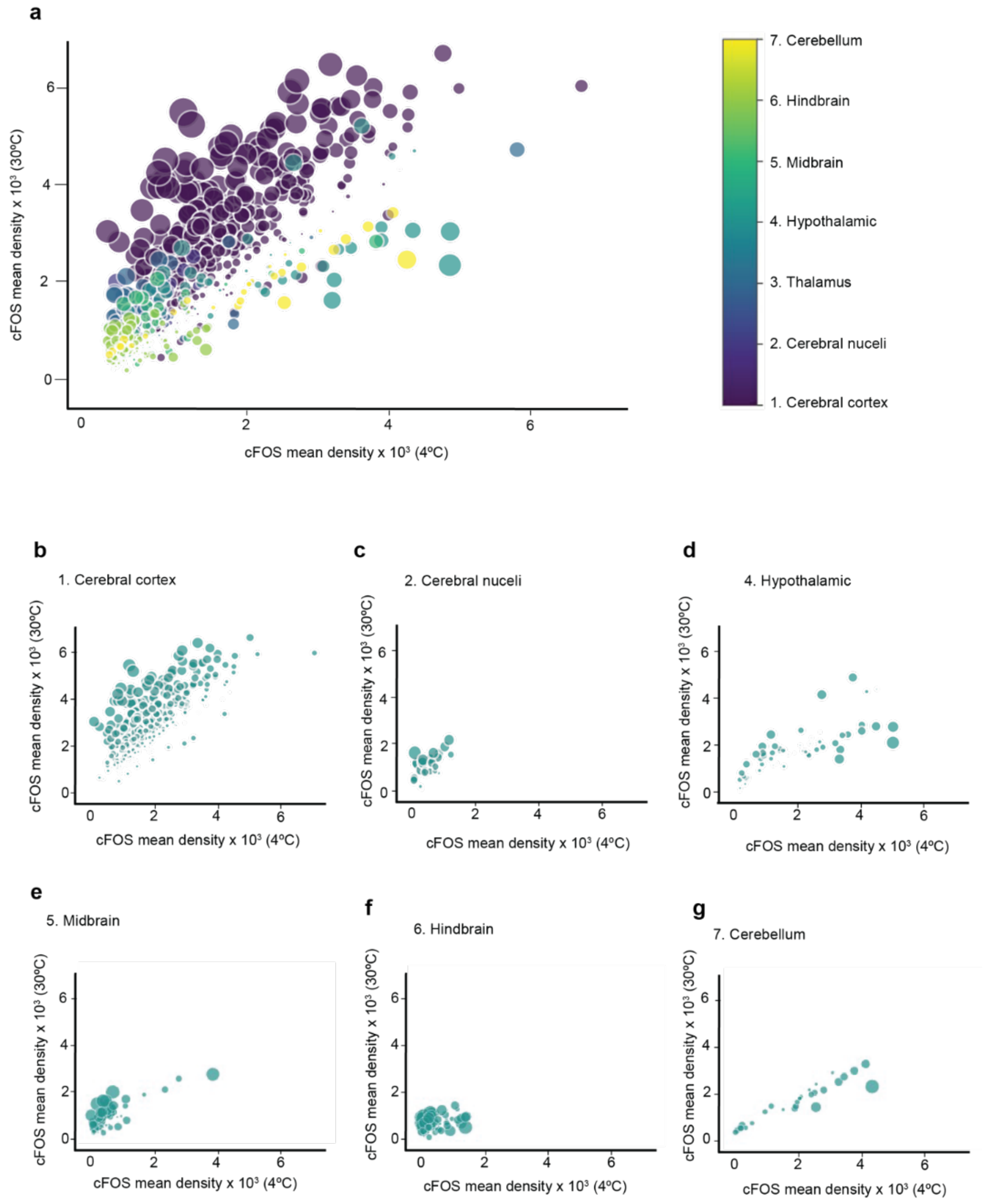


**Extended Data Fig. 3| Whole-brain screening of mice in non-CIEC 30°C or CIEC 4°C state. a-g,** Brains were harvested from mice undergoing CIEC (6 hrs at cold 4°C) or non-CIEC (6 hrs at thermoneutral 30°C). n=4 animals per group. Whole-brain imaging and cFos mapping data for the entire brain in **a** and individual regions of the cerebral cortex in **b**, cerebral nuclei **c**, hypothalamus **d**, midbrain **e**, hindbrain **f**, and cerebellum **g**. Each dot represents an annotated brain region based on the Allen Brain Atlas. The size of the dots represents the mean differences between warm and cold conditions in each region. Also see supplementary table 1 for all regions. See also Supplementary Table 1.

**Extended Data Figure 4**

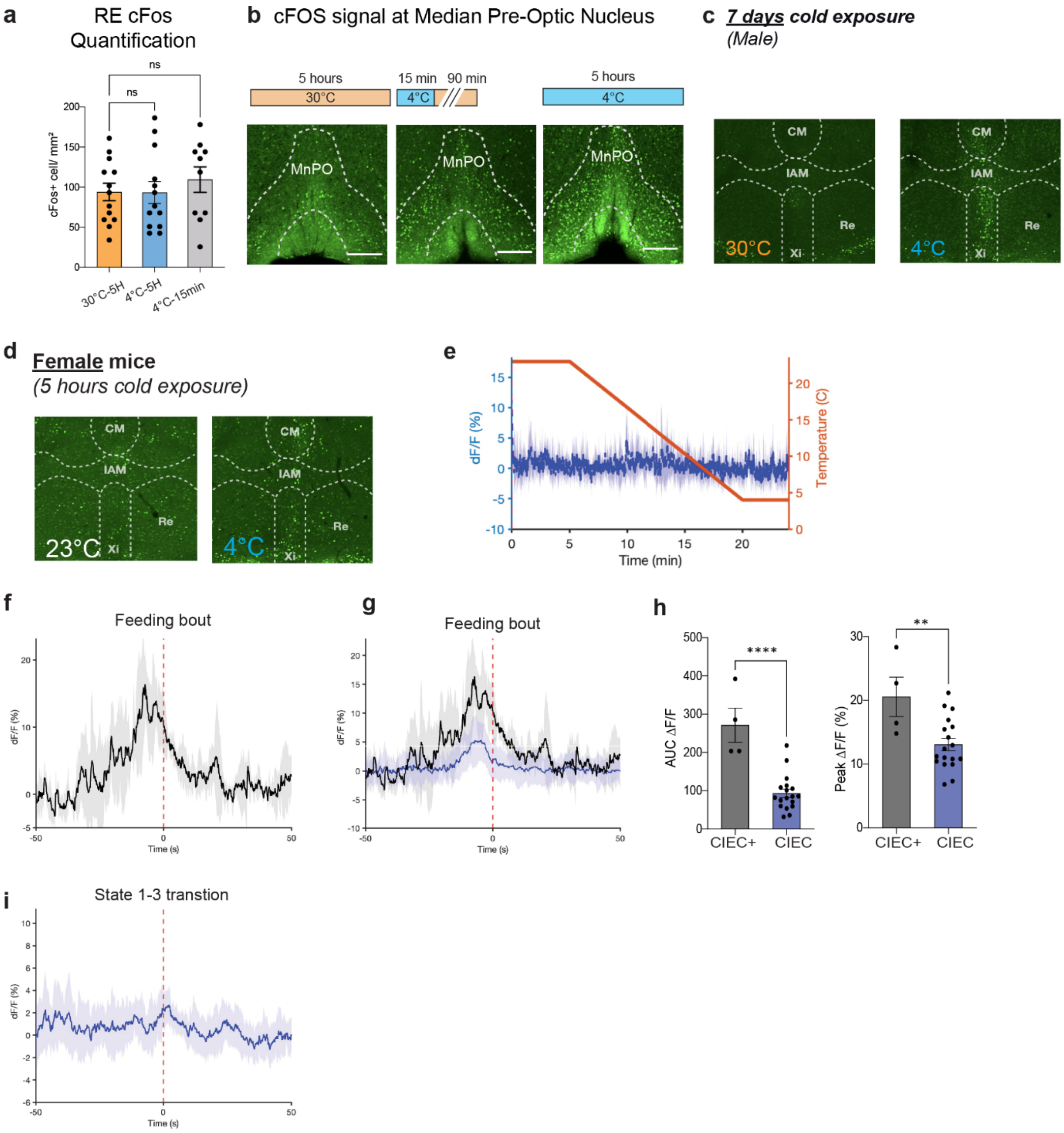


**Extended Data Fig. 4| Activity for Xi neurons during cold sensation vs. CIEC and HMM state transitions. a,** cfos quantification of Re region from main Fig. 2b. **b,** representative images of MnPO (at Bregma 0.14) for same experiment as shown in main Fig. 2a-c. **c,** Representative Xi images from mice after 7 days of cold exposure (n=4 for RT and cold). **d,** Representative Xi c-Fos images from female mice after 5 hours of cold exposure. (n=2 for RT, n=3 for cold). Scale bar: 500 µm. **e,** Average calcium signal from Xi neurons while the ambient temperature ramped down from 23°C to 4°C. (averaged from n=3 mice). **f-g,** Xi calcium signal for mice undergoing CIEC or a CIEC+ state (4°C without food for 3 hrs to generate exacerbated cold-induced energy compensation), dotted red line indicates a single feeding bout. (n=4 mice). **h,** Quantification of AUC delta F/F and peak delta F/F. Data are mean ± SEM. ****$P < 0.0001$, **$P < 0.01$ using an unpaired t-test. **I,** Averaged calcium signal from Xi neurons from mouse undergoing CIEC, averaged from 11 events from 3 different mice. The red dotted line is the state transition time point calculated using HMM.

### Extended Data Figure 5

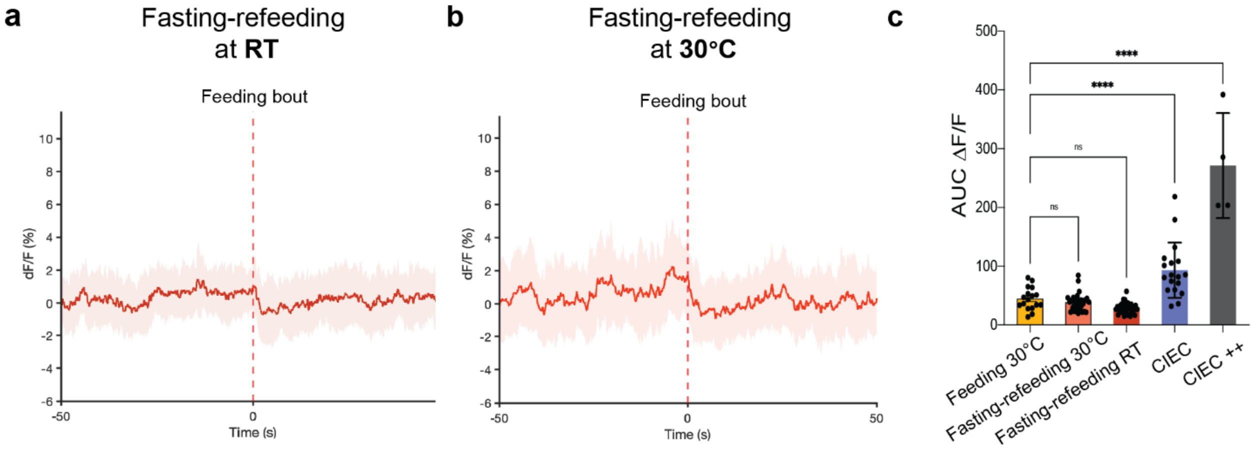


**Extended Data Fig. 5| Specificity of Xi neuronal activity during different energy deficit states. a-b,** Fiber photometry signal of AAV-GCaMP6m expressing vMT/Xi neurons after overnight fasting during refeeding at room temperature (**a**) or at 30°C **(b**), shown as the average of 35 events from 5 different mice for (**a**) and 31 events from 5 different mice for (**b**). **c,** Bar graph showing the area under the curve (AUC) dF/F (-20s to 10s) for **a,b** and main Fig. 2f and g. Data are mean ± SEM. ****P < 0.0001, ns=0.8572 for comparison between 30°C feeding and fasting-refeeding at 30°C, ns=0.1678 for comparison between 30°C feeding and fasting-refeeding at 23°C using a one-way ANOVA with Dunnett's multiple comparison test.

**Extended Data Figure 6**

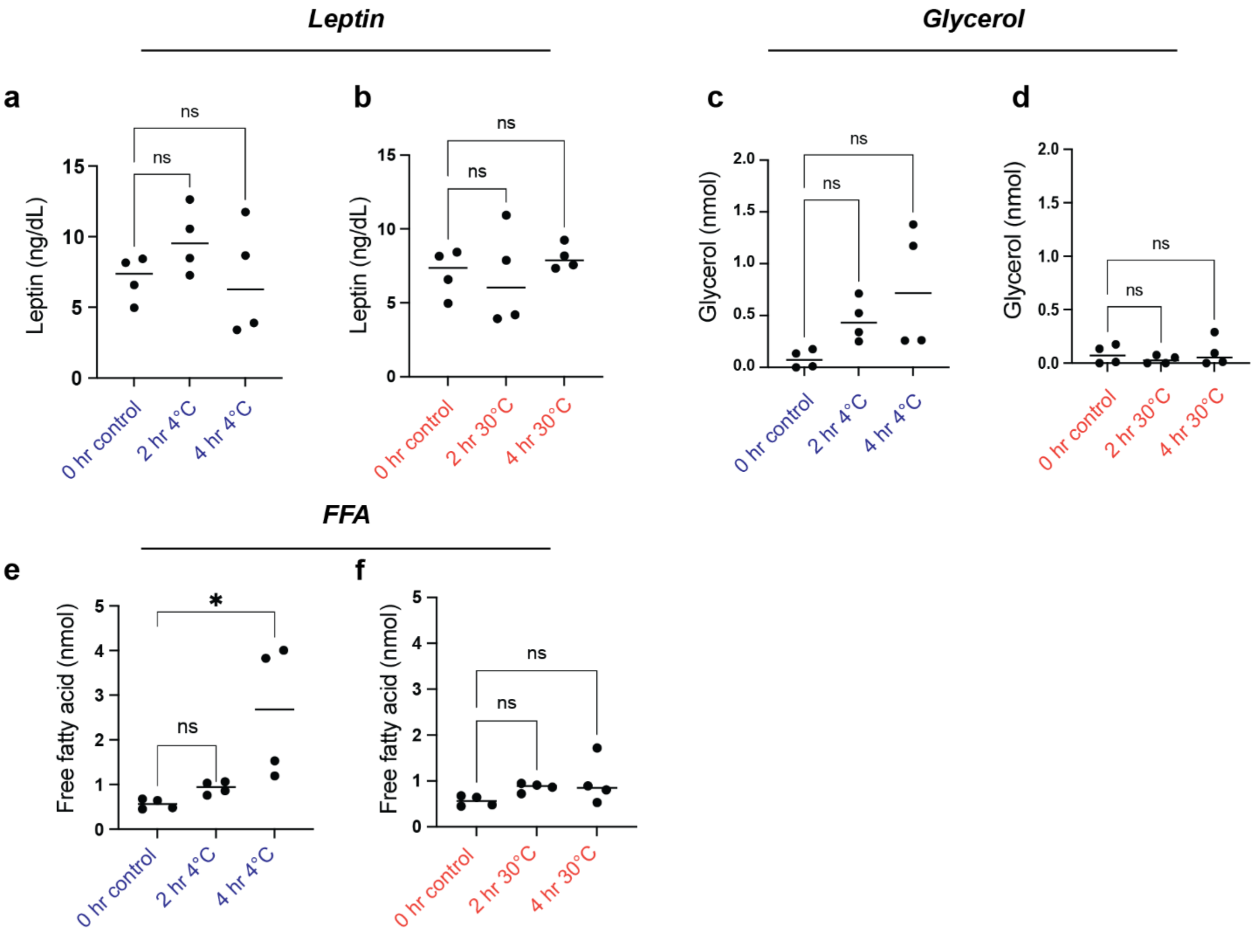


**Extended Data Fig. 6| Hormonal and metabolic profiling of mice undergoing CIEC vs mice exposed to thermoneutral conditions. a-b,** Quantification of circulating leptin levels in mice after being in cold **(a)** or at 30°C (thermoneutral) **(b)** for given amount of time. **c-f,** Quantification of plasma glycerol (**c** and **d**) and free fatty acids (**e** and **f**) for mice in cold or or at 30°C (thermoneutral). Data are mean ± SEM. *P = 0.0143 for FFA in cold using a one-way ANOVA with Dunnett's multiple comparison test.

### Extended Data Figure 7

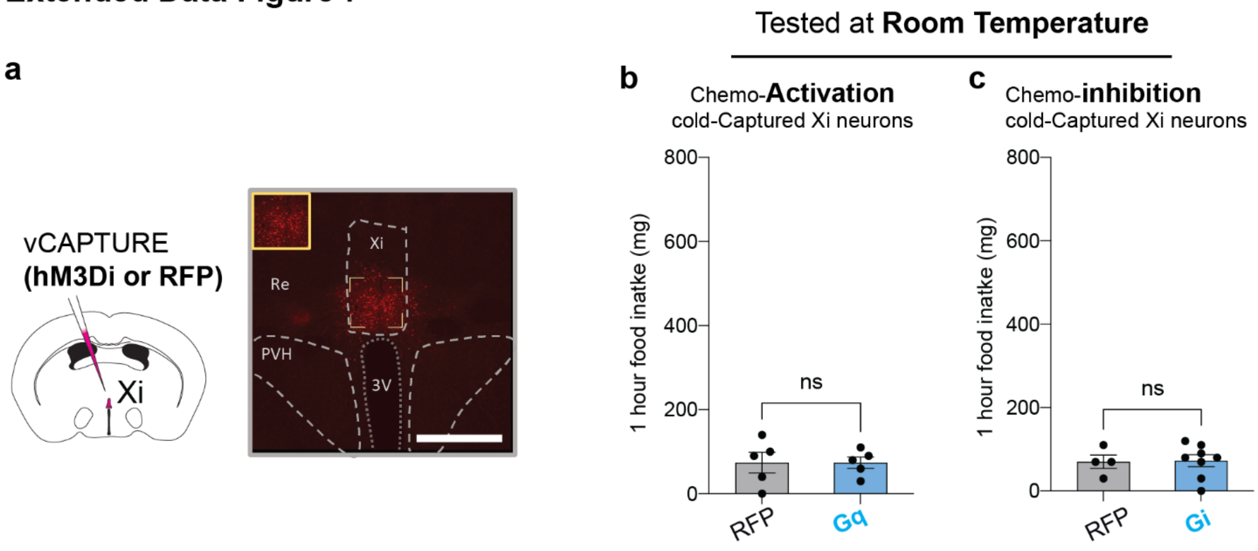


**Extended Data Fig. 7| Reactivation of cold- vCAPTURE DREADD Xi neurons at room temperature. a,** Schematic and representative histology section from vCAPTURE DREADD Gi mouse. Scale bar: 500 µm. **b,** Food intake for cold-vCAPTURED Xi Gq activated at 23°C (n=5 for RFP and Gq). **c,** Food intake data for cold-vCAPTURED Xi Gi at 23°C (n=4 for RFP and n=8 for Gi). Data are mean ± SEM. ns >0.99 for Gq, ns =0.9163 for Gi food intake at 23°C using an unpaired t-test.

**Extended Data Figure 8**

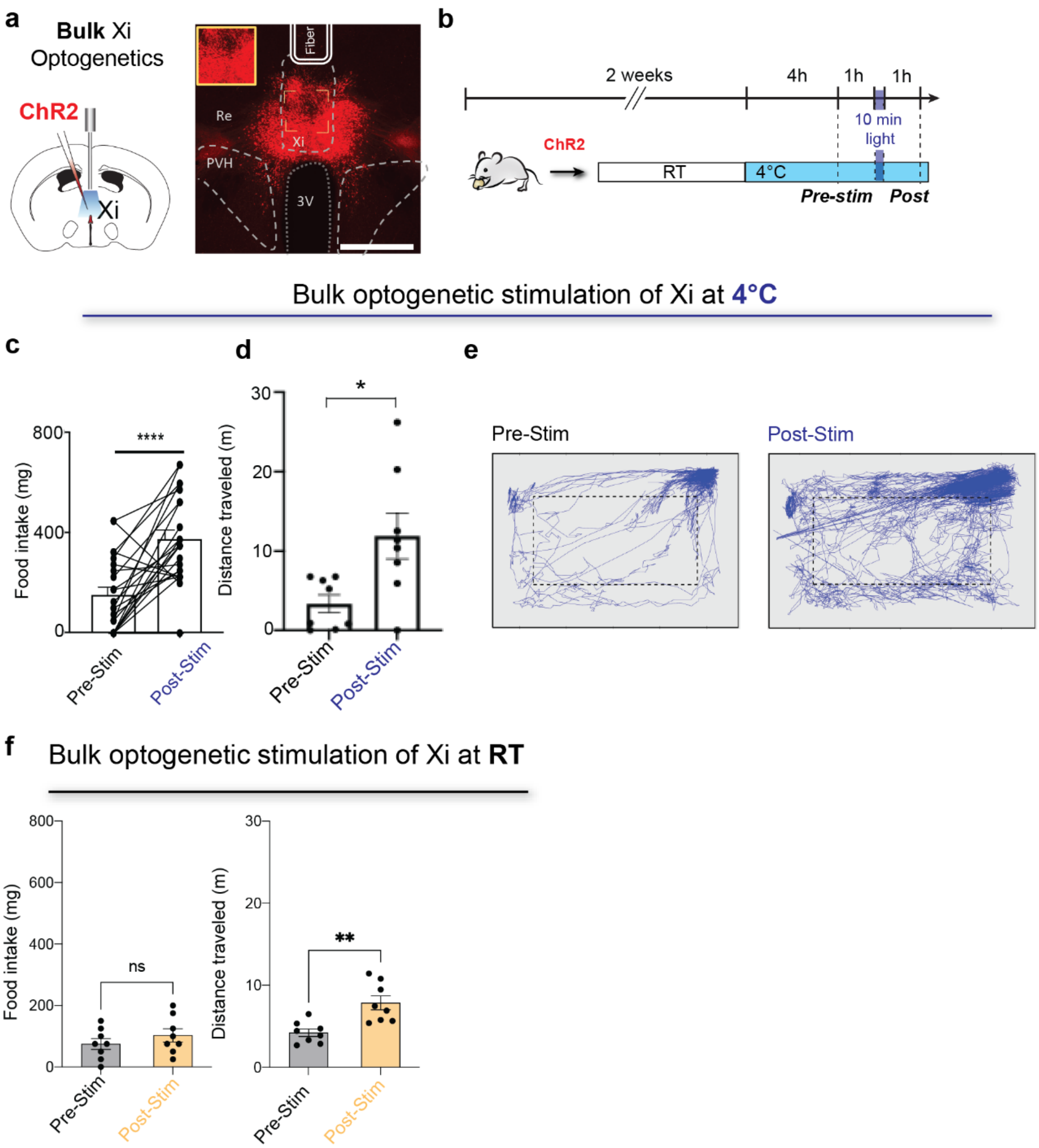


**Extended Data Fig. 8| Constitutive optogenetic activation of Xi neurons during CIEC. a,** Schematic and representative histology section from a mouse injected with constitutive (hSyn-ChR2) at the Xi. Scale bar: 500 μm. **b,** Experimental design for testing the effect of bulk activation of Xi and surrounding neurons on CIEC-associated feeding. **c-e,** Food intake pre- and post-stimulation (n=8 mice, N=16 times)(**c**) and **d-e** physical activity (measured as total distance traveled) pre- and post-stimulation (**d**) and individual traces of physical movement of a single animal (**e**). **f,** Food intake and physical activity of animals expressing constitutive ChR2 in the Xi at RT (n=8 animals) Data are mean ± SEM. ****P < 0.0001, **P=0.0096, *P < 0.01 using a paired t-test.

**Extended Data Figure 9**

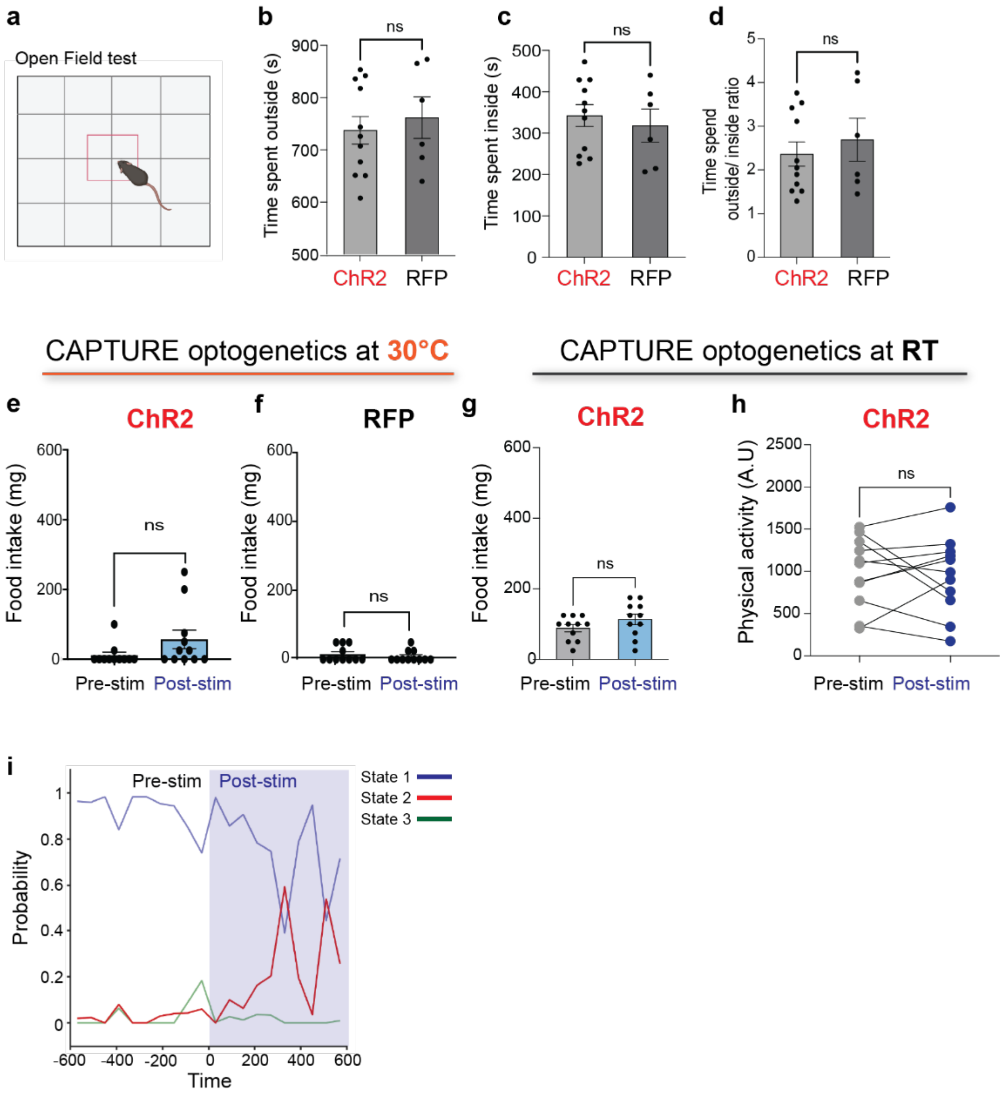


**Extended Data Fig. 9| vCAPTURE Xi-CIEC optogenetics, open field test, and HMM state transition. a,** Schematic of the open field test. **b-d,** Quantification of time spent outside (**b**) or time spent inside (**c**) (or the ratio thereof (**d**)) following activation of vCAPTUREd Xi$^{\text{CIEC}}$ neurons in ChR2 or RFP control mice (n=11 for ChR2 and n=6 for RFP). **e-f,** Bar graph showing the difference in food intake at thermoneutral temperature pre- and post-laser stimulation for ChR2 mice in **e** and RFP mice in **f.** Data are mean ± SEM. ns= 0.1501 for **e** and ns=0.4650 for **f** using paired t-test. **g-h,** Food intake (**g**) and physical activity (**h**) for vCAPTURED Xi$^{\textbf{CIEC}}$ mice stimulated at room temperature (n=11 mice). Data are mean ± SEM. ns= 0.1530 for **g** and ns=0.7614 for **h** using paired t-test. **i,** Continuous state probability calculated for ChR2 mouse, pre-stimulation (white background) or post-stimulation (blue background) while undergoing CIEC (n=7 mice).

**Extended Data Figure 10**

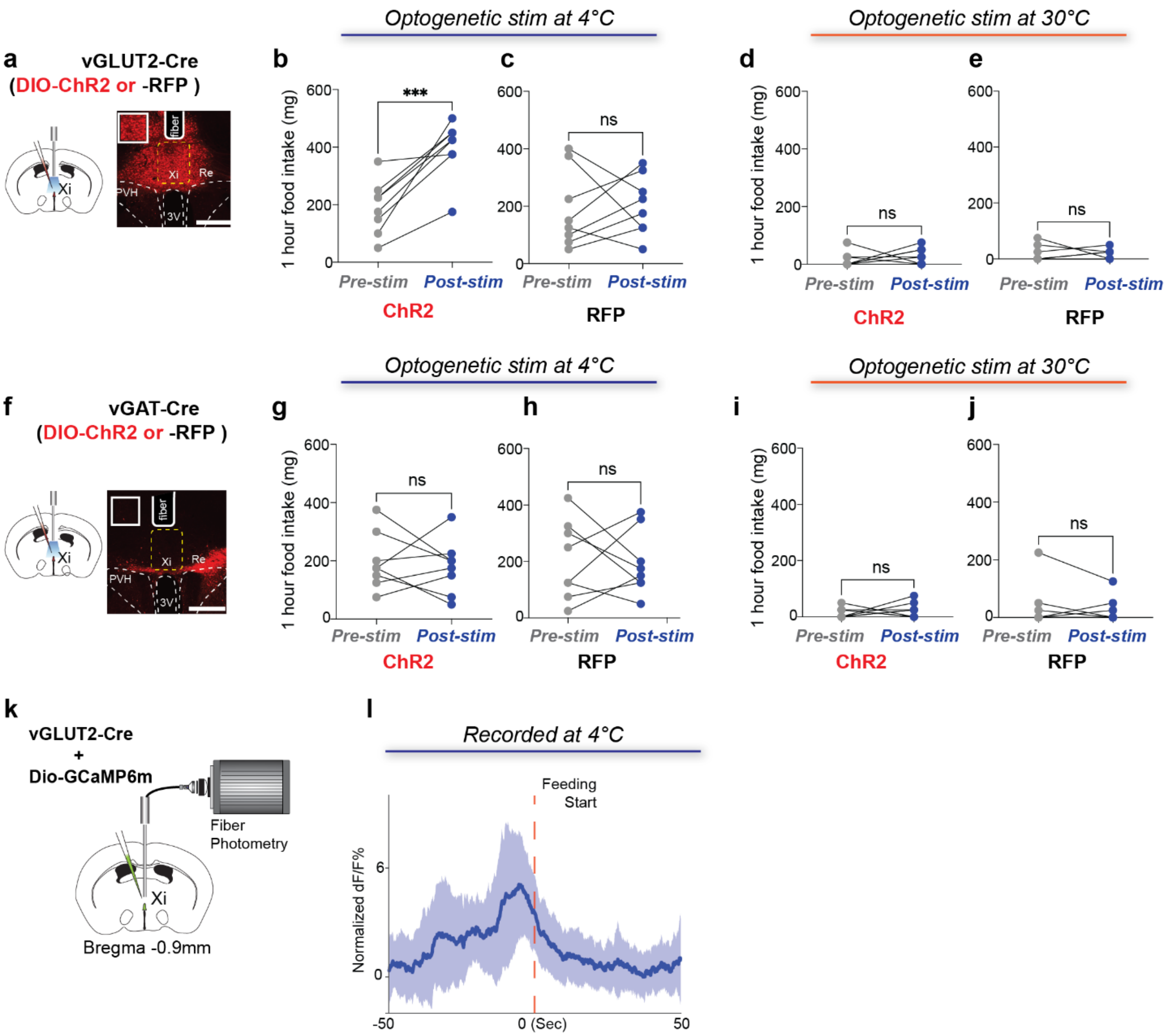


**Extended Data Fig. 10| Xi$^{vGLUT2}$ neurons mediate CIEC-associated behavior.**
**a,** Schematic and representative histology showing injection of a viral AAV-DIO-ChR2 and optogenetic implant at the Xi (solid white line indicates fiber track) in vGLUT2 animals. Scale bar: 500 µm. **b-c,** Food intake in cold pre- and post-stimulation for vGLUT2-ChR2 mice (**b**) (n=8 mice) and for vGLUT2-RFP control mice (**c**) (n=8 mice). **d-e,** Food intake for vGLUT2-ChR2 mice (**d**) and vGLUT2-RFP mice (**e**) pre- and post-stimulation under thermoneutral conditions. **f,** Schematic and histology of vGAT2 animals. Scale bar: 500 µm. **g-j**, Food intake in cold pre- and post-stimulation (**g-h**), and under thermoneutral conditions (**i-j**). Data are mean ± SEM. ***$P < 0.009$ for **b**, ns= 0.7849 for **c**, ns= 0.4869 for **d**, ns= 0.8018 for **e**, ns= 0.6682 for **g**, ns= 0.8383 for **h**, ns= 0.5490 for **i**, ns= 0.47 for **j** by paired t-test. **k**, Schematic of fiber photometry setup for

recording from vGLUT2-Xi injected with Dio-GCamp6m at the Xi in vGLUT2 animals. **I,** Fiber photometry signal of AAV-Dio-GCaMP6m expressing in vGLUT2-Xi neurons in the cold, shown as the average of 15 events from 4 different mice.

**Extended Data Figure 11**

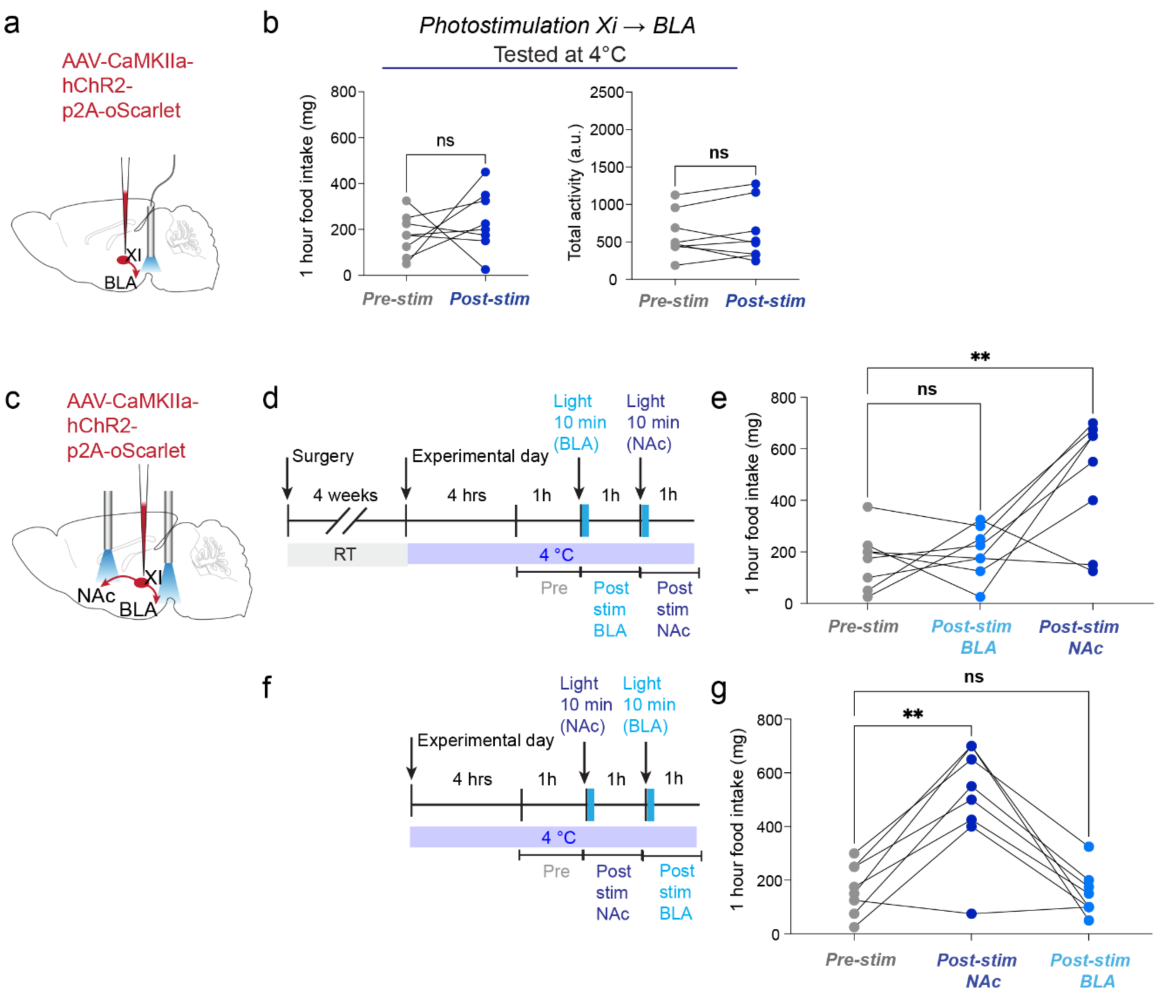


**Extended Data Fig. 11| Xi-NAc but not Xi-BLA regulate CIEC-associated feeding.** **a,** Schematic of viral AAV-ChR2 injection at the Xi and optogenetic implant above the BLA in WT animals. **b,** Food intake and physical activity in cold pre- and post-laser stimulation for Xi-BLA mice (n=8 mice)**. c**, Schematic of AAV-ChR2 injection at the Xi and optogenetic implant above both the BLA and NAc in the same animal. **d-e,** Timeline (**d**) and food intake data (**e**) for the experiment in which the BLA was stimulated first and then the NAc. Food intake is plotted for pre-stimulation, post-stim for the BLA, and post-stim for the NAc. **f-g,** Timeline (**f**) and food intake data (**g**) for experiment in which the NAc was stimulated before the BLA. Data are mean ± SEM. **P < 0.0063, ns=0.8083 for **e,** **P < 0.0028, ns=0.7924 for **g** using a one-way ANOVA with Dunnett's multiple comparison test.
